# Self-reported Health is Related to Body Height and Waist Circumference in Rural Indigenous and Urbanised Latin-American Populations

**DOI:** 10.1101/562942

**Authors:** Juan David Leongoméz, Oscar R. Sánchez, Milena Vásquez-Amézquita, Eugenio Valderrama, Andrés Castellanos-Chacón, Lina Morales-Sánchez, Javier Nieto, Isaac González-Santoyo

## Abstract

Body height is a life-history component. It involves important costs for its expression and maintenance, which may originate trade-offs on other costly components such as reproduction or immunity. Although previous evidence has supported the idea that human height could be a sexually selected trait, the explanatory mechanisms that underlie this selection are poorly understood. Despite extensive studies on the association between height and attractiveness, the role of immunity in linking this relation is scarcely studied, particularly in non-Western populations. Here, we tested whether human height is related to health measured by self-perception, and relevant nutritional and health anthropometric indicators in three Latin-American populations that widely differ in socioeconomic and ecological conditions: two urbanised populations from Bogota (Colombia) and Mexico City (Mexico), and one isolated indigenous population (Me’Phaa, Mexico). Results showed that self-reported health is best predicted by an interaction between height and waist circumference, and the costs associated with large waist circumference are height-dependent, affecting taller people more than shorter individuals. If health and genetic quality cues play an important role in human mate-choice, and height and waist interact to signal health, its evolutionary consequences, including cognitive and behavioural effects, should be addressed in future research.

## Introduction

In modern Western societies, it has been seen that women usually prefer men who are significantly taller than average^1–3^, while men are more tolerant in choosing women who are taller or shorter than average ^4^. This is consistent with the idea that male height can be adaptive^5^ and sexual selection favours taller men, possibly because height may represent a honest signal of individual quality, providing hereditary advantages, such as genetic quality for the offspring^6, 7^, or direct benefits, provisioning resources and protection for women and their children^8^. Following these last possible benefits, height has been also proposed as an indicator of resource holding potential (RHP), in terms of social dominance and deference ^9, 10^ and socioeconomic status^6, 11^.

This idea is supported by evidence that the male height is directly correlated with reproductive success, which is not applicable to women, suggesting an unrestricted directional selection that favours very tall men but not to very tall women^12^. In fact, it has been reported that taller men (but not extremely tall men) are more likely to find a long-term partner and have several different long-term partners^13^, while the maximum reproductive success of women is below the female average height^14^. Furthermore, heterosexual men and women tend to adjust their preferred height of hypothetical partners according to their own stature^15^. In general, heterosexual men and women prefer couples in which the man is taller than the woman, and women show a preference for facial cues that denote a taller man^16^.

Although previous evidence has supported the idea that human height could be a sexually selected trait, little is known regarding whether this role is based on the honest signalling of individual quality^6^. To test this idea, it is important to consider that height needs to face a trade-off with other life history components^17^, such as reproduction^18^ or immunity^19^, and its expression should involve certain costs that not all individuals could equally afford, such that there would be an important phenotypic variation on this trait. Both aspects are present in human height. First, growth in body height is a life-history component^1, 20^ that involves important costs for its expression and maintenance. The costs in height can be measured in terms of survival and physiological expenditure^19^. For example, according to the Hayflick limit theory of ageing^21^, our cells have a limited number of cell replications available in a lifetime. A minimal increment in body height involves more cells, maybe trillions, and large numbers of cell replications. These large numbers of cell replications demand a large pool of proteins to maintain taller, larger bodies^19^, which together, with an increase of free radicals generated by normal cellular metabolism, may lead to a greater likelihood of DNA damage^22^, thus increasing the incidence of cancer and reducing longevity^19^.

Secondly, reproduction and immunity could face a trade-off with height because the effect of sex hormones on these life history components; the trade-off with reproduction occurs when the notorious increment of sex steroids, particularly testosterone and oestrogens, induce an accelerated growth period in puberty for both sexes^23^, but also a reallocation of physiological resources for reproduction posterior to this period (i.e. spermatogenesis, follicular maturation, etc.), which results in a growth cessation. Both steroids stimulate mineral deposition in the growth plates at the ends of the long bones, thus terminating cell proliferation and resulting in the fusion of the growth plates to the shaft of the bone^24, 25^.

In turn, the increment of sexual steroids at sexual maturity trigger another trade-off for individuals that is particularly associated with the effect of testosterone on immunity^26^. Usually, testosterone exerts suppressive effects on several, but not all, immune components^27^. For example, it may negatively affect the activity and cellular proliferation of several adaptive and innate immune responses, such as Macrophages, Natural Killers, production of cytokines^28^ and helper T cells and lymphocyte activation (i.e. Th2 and Th17^29^). In consequence, it has been documented that testosterone may influence general health patterns, in relation to the severity of certain infections such as malaria, leishmaniasis, amoebiasis^30^, and tuberculosis^31, 32^

Therefore, as a consequence of these life-history trade-offs, height could be considered as a reliable indicator of an individual’s condition in terms of (1) the amount and quality of nutritional resources acquired until sexual maturity, (2) the RHP to obtain resources for the somatic maintenance in the adult stage, and (3) the general health condition, given the possibly costly immunosuppressive effect of testosterone. Thus, height can be used for potential partners to receive information about the quality of potential mates; only high-quality individuals could afford to allocate resources to this attractive secondary sexual trait^33^, which would result in an increased sexual preference towards taller individuals.

Despite extensive studies on the association between height and attractiveness, the role of height as signal of biological quality has been largely studied but results are still controversial. For example, one aspect that has been particularly studied is the relationship between height and various indicators of health. Generally, height is positively related to measures of health in men, such as coronary heart disease morbidity and mortality^34^, limiting long-standing illness^6^ and perceived health^6, 35^. Nevertheless, the implementation sex and of other indicators of health has led to conflicting results, demonstrating the complexity of the question. For instance, mortality by cancer diseases has been associated more to taller than shorter people of both sexes^36^, and height somewhat predicts general health in women but following a curvilinear trend^6^.

Moreover, most studies have been done using high-income developed populations such as Western, Educated, Industrialised, Rich and Democratic (WEIRD) societies^37^, which has led to a lack of information of what is occurring in other populations with important socio-ecological differences^35^. These ecological pressures are important because although genetic allelic expression could be the main factor that determines individual height differences^35^, height is also the most sensible human anatomical feature that responds to environmental and socioeconomic conditions^18, 38^. For instance, variation in height across social classes is known to be greater in poorer countries^39^ but is much reduced in countries with higher standards of living^40^. Economic inequality not only affects the population’s nutritional patterns, which are especially important during childhood to establish adult height, but also the presence of infectious diseases^41^. Childhood disease is known to adversely affect growth. For instance, mounting an immune response to fight against the infection requires concomitant increases in metabolic rate, which could affect the net nutrition, and hence reduces productivity. Disease also prevents food intake, impairs nutrient absorption and causes nutrient loss^42, 43^. Therefore, compared with high-income and developed populations, habitants from locations with stronger ecological pressures imposed by pathogens or greater nutritional deficiencies would face greater costs to robustly express this trait, thereby showing stronger sexual selective pressure over height, as it signals growth rates, life-history trajectories and health status more accurately. This phenotypic variation is described as developmental plasticity, which is a part of the phenotypic plasticity related to growth and development, in response to social, nutritional and demographic conditions, among others^44^. During the last century, given a general improvement in nutrition, human height has steadily increased across the globe^45^, but the level of dimorphism in favour of men is maintained.

Colombia and Mexico are two of the most socioeconomically heterogeneous countries in the world with a high Human Development Index^46^. Colombia and Mexico attain respective scores of 68 and 66 in the Healthcare Access and Quality Index^47^, indicating that the populations are in relatively good health compared to global standards. Yet, Colombia and Mexico have GINI coefficients of 50.8 and 43.4, respectively, making them the 12^th^ and 43^rd^ most unequal countries in the world (GINI index – World Bank estimate; https://data.worldbank.org/indicator/SI.POV.GINI). These national-level statistics, however, hide important within-country differences. In particular, Latin-American people in rural areas tend to be poorer and have less access to basic services such as health and education than people in urban areas.

According to data from the World Bank and the Colombian National Administrative Department of Statistics, in 2017 Colombia was the second most unequal country in Latin-America after Brazil. In rural areas, 36% of people were living in poverty and 15.4% in extreme poverty, while in urban areas, these values were only 15.7% and 2.7%, respectively^48^.

In addition to rural communities, in Latin-America indigenous people tend to have high rates of poverty and extreme poverty^49^, and poorer health^50^, which is less susceptible to improvement by national income growth^51^. In Mexico, there are at least 56 independent indigenous peoples whose lifestyle practices differ in varying degrees from the typical ‘urbanised’ lifestyle. Among these groups, the Me’Phaa people, from an isolated region known as ‘*Montaña Alta*’ of the state of Guerrero, is one of the groups whose lifestyle most dramatically differs from the typical Westernised lifestyle of more urbanised areas^52^. Me’Phaa communities are small groups of indigenous people, composed of 50 to 80 families, each with five to ten family members. Most communities are based largely on subsistence farming of legumes such as beans and lentils, and the only grain cultivated is corn. Animal protein is acquired by hunting and raising some fowls, and meat is only consumed during special occasions but not as part of the daily diet. There is almost no access to allopathic medications, and there is no health service, plumbing or water purification system. Water for washing and drinking is obtained from small wells. Most of the Me’Phaa speak only their native language^53^. In consequence, these communities have the lowest income and economic development in the country, and the highest child morbidity and mortality due to chronic infectious diseases^52^.

These three Latin-American populations can provide an interesting indication about how the regional socioeconomic conditions and intensity of ecological pressures by pathogens may modulate the function of height as an informative sexually selected trait of health and individual condition in each sex. Therefore, the aim of the present study was to evaluate whether human height is related to health measured by self-perception, and relevant nutritional and health anthropometric indicators in three Latin-American populations that widely differ in socioeconomic and ecological conditions: two urbanised populations from Bogota (Colombia) and Mexico City (Mexico), and one isolated indigenous population (MéPhaa, Mexico). In addition, given the possible immunological effects of testosterone, and that men present higher levels than women, we predicted this relation to be different between sexes in all studied populations. Therefore, we propose that height would be a stronger signal of self-reported health condition in men compared to women.

## Methods

All procedures for testing and recruitment were approved by Universidad El Bosque Institutional Committee on Research Ethics (PCI.2017-9444) and National Autonomous University of Mexico Committee on Research Ethics (FPSI/CE/01/2016), and run in accordance with the ethical principles and guidelines of the Colombian College of Psychologists (COLPSIC) and the Official Mexican Law (NOM-012-SSA3-2012). All participants read and signed a written informed consent.

### Participants

A total of 477 adults (238 women and 239 men) participated in this study. They were from three different samples: (1) Mexican indigenous population, (2) Mexican urban population and (3) Colombian urban population. In Mexico, Me’Phaa indigenous participants from ‘La *Montaña Alta*’ were recruited and participated in this study between January and March 2017, while data from participants from Mexico City was collected between May and June 2017. In Colombia, data collection was carried out between October 2018 and May 2019.

The first sample consisted of 63 subjects (mean age ± standard deviation [SD] = 33.63 ± 9.69 years old) from the small Me’Phaa community – ‘*Plan de Gatica*’ from a region known as ‘*Montaña Alta*’ of the state of Guerrero in Southwest Mexico. In this sample, 24 participants were women (33.46 ± 8.61 years old) and 39 participants were men (33.74 ± 10.41 years old), who participated in a larger study on immunocompetence. Both sexes were aged above 18 years old. In Mexico, people above 18 years old are considered adults. All measurements were collected in the participants’ own community. Me’Phaa communities are about 20 km apart, and it takes about three hours of travel on rural dirt roads to reach the nearest large town, about 80 km away. Mexico City is about 850 km away, and the trip takes about twelve hours by road. This community has the lowest income in Mexico, the highest index of child morbidity and mortality by gastrointestinal and respiratory diseases (children aged 0 to 8 years had the highest vulnerability and death risk^52^), and the lowest access to health services. These conditions were recorded in the National Health Information System 2016^52^.

The second sample consisted of 60 subjects of over 18 years old (30.27 ± 8.56 years old) from the general community in Mexico City, of whom 30 were women (37.47 ± 5.61 years old) and 30 were men (23.07 ± 3.22 years old). Finally, the third sample consisted of 354 undergraduate students with ages ranging from 18 to 30 years old (20.39 ± 2.10 years old), 184 were women (20.16 ± 2.08 years old), and 170 were men (20.64 ± 2.10 years old) from Bogota, Colombia. All urban participants were recruited through public advertisements.

Participants from both urban population samples were taking part in two separate, larger studies in each country. In Colombia, all data were collected in the morning, between 7 and 11 am, because saliva samples (for hormonal analysis), as well as voice recordings, body odour samples, and facial photographs were also collected as part of a separate project. Additionally, women in the Colombian and Mexican samples were not hormonal contraception users, and all data were collected within the first three days of their menses.

Participants who were under allopathic treatment and hormonal contraception users from both countries were excluded from data collection. All participants completed a sociodemographic data questionnaire, which included medical and psychiatric history. No women were users of hormonal contraception. Although no participant reported any endocrinological or chronic disease, these health issues were also considered as exclusion criteria.

Given that the indigenous community of ‘*Plan de Gatica*’ consists of 60–80 families, each with five to seven members, the final sample for this study could be considered as semi-representative of a larger Me’Phaa population inhabiting in the same community. Nevertheless, the total population of Me’Phaa people inhabiting the ‘*Montaña Alta*’ is comprised of 20–30 communities with almost the same number of families as ‘*Plan de Gatica*’. Therefore, it is important to mention that our sample size cannot be considered representative of the total Me’Phaa people inhabiting the ‘*Montaña Alta*’ region, but from the specific ‘*Plan de Gatica*’ community. Similar condition occurs for participants from the Mexico City and Bogota samples. These participants were recruited at the National Autonomous University of México and Universidad El Bosque campuses, respectively. Therefore, these samples are comprised mostly of bachelor and graduate students, and cannot be considered as representative of a large population of the whole city, which is comprised of about 12 million adult persons in Mexico and about 5 million adults in Bogota.

### Procedure

All participants signed the informed consent and completed the health and background questionnaires. For participants from the indigenous population, the whole procedure was carried out within their own communities, and participants from the Mexican and Colombian urban population attended a laboratory at either the National Autonomous University of México or Universidad El Bosque respectively, on individual appointments.

Participants from Mexico City and Bogota were recruited through public advertisements on social media and poster boards located along the central campus of the National Autonomous University and Universidad El Bosque. While in Mexico City, participants received either one partial course credit or a payment equivalent to $5 dollars as compensation for their participation, all participants in Bogota were given academic credits for their participation.

For the indigenous groups, recruitment was done through the Xuajin Me’Phaa non-governmental organisation, which is dedicated to the social, environmental and economic development for the indigenous communities of the region (see video from this organisation, http://youtu.be/In4b9_Ek78o). Xuajin Me’Phaa has extensive experience in community-based fieldwork and has built a close working relationship with the community authorities. The trust and familiarity with the community customs and protocols have previously led to successful academic collaborations^53, 54^. Therefore, Xuajin Me’Phaa served as a liaison between the Mexican research group the and communities for the present study, offering mainly two important factors in data collection: the informed consent of community members and participants, and two trained interpreters of Me’Phaa and Spanish language of both sexes.

First, participants were asked to complete the health and sociodemographic data questionnaires. Subsequently, the anthropometric measurements were taken.

#### Self-reported health

In order to obtain a standardised value of self-perception of health, we implemented in all three populations the Short Form (36) health survey (SF-36; RAND Corp.; https://www.rand.org/health-care/surveys_tools/mos/36-item-short-form/survey-instrument.html). The SF-36 produces eight dimensions, of which we only used the las one (8) General health. Each factor is calculated by averaging the recoded scores of individual items: (1) Physical functioning (items 3 to 12), (2) Role limitations due to physical health (items 13 to 16), (3) Role limitations due to emotional problems (items 17 to 19), (4) Energy/fatigue (items 23, 27, 29 and 31), (5) Emotional well-being (items 24, 25, 26, 28 and 30), (6) Social functioning (items 20 and 32), (7) Pain (items 21 and 22) and (8) General health (items 1, 33, 34, 35 and 36).

The interpreters provided by the Xuajin Me’Phaa organisation administered the SF-36 Health survey in Me’Phaa language. Interpreters used Spanish as the second language and are thoroughly proficient in speaking and reading Spanish. We used the validated SF-36 survey for urban and rural Mexican populations^55^ for interpreters to translate Spanish to Me’Phaa language. Given the ethnical customs of Me’Phaa culture, the participants were always interviewed by an interpreter of the same sex to avoid bias in participant responses; for instance, men were interviewed by a male interpreter and women by a female interpreter. The same interpreter interviewed all participants of his/her corresponding sex.

For the present study, both urban and indigenous participants only answered items corresponding to the dimension defined as general health (i.e. Item numbers 1, 33, 34, 35 and 36), except for item 35. This item informs about the expectation for future health. Since the grammatical compositions of Me’Phaa language do not consider ‘infinitive’ and ‘future’ as verbal tenses^56^, an interpretation of this question was not possible for the Me’Phaa people, therefore, this item was excluded.

In Colombia, we used a Spanish version of the SF-36 questionnaire ^57^, that was previously validated in the same country^58^.

To obtain the self-reported health rate, all items were recoded following the instructions on how to score SF-36^57^. We calculated the final factor by averaging the recoded items. To make this data compatible with the Mexican database, item 35 was excluded because it cannot be answered by the Mexican Indigenous population, and the general health dimension was calculated by averaging only items 1, 33, 34 and 36.

#### Anthropometric measurements

All anthropometric measurements were measured thrice and subsequently averaged to obtain the mean value (for agreement statistics between the three measurements of each characteristic, see section 1.3 in the Supplementary Material). All participants wore light clothing and had their shoes removed. The same observer repeated the measurements thrice.

We measured the body height in cm, to the nearest mm, by using a 220 cm Zaude stadiometer, with the participant’s head aligned according to the Frankfurt horizontal plane, and feet together against the wall.

Anthropomorphic measurements also included waist circumference (cm), weight (kg), fat percentage, visceral fat level, muscle percentage and body mass index (BMI). The waist circumference was measured midway between the lowest rib and the iliac crest in cm by using a flexible tape and was recorded to the nearest mm. These anthropomorphic measurements have been used as an accurate index of nutritional status and health, especially waist circumference. Metabolic syndrome is associated with visceral adiposity, blood lipid disorders, inflammation, insulin resistance or full-blown diabetes and increased risk of developing cardiovascular disease^59–61^, amongst Latin-American populations^62^. Waist circumference has been proposed as a crude anthropometric correlate of abdominal and visceral adiposity, and it is the simplest and accurate screening variable used to identify people with the features of metabolic syndrome^63, 64^. Hence, in the presence of the clinical criteria of metabolic syndrome, increased waist circumference provides relevant pathophysiological information insofar as it defines the prevalent form of the syndrome resulting from abdominal obesity^60^.

Weight, fat percentage, visceral fat level, muscle percentage and BMI were obtained using an Omron Healthcare HBF-510 body composition analyser, which was calibrated before each participant’s measurements were obtained.

### Statistical analysis

We used linear models (LM) to test the association between height and self-reported health. The dependent variable in this model was the health factor and predictor variables included participant sex, age, sample (Bogota, Mexico City, Me’Phaa), height and waist and anthropometric measurements (hip, weight, fat percentage, BMI and muscle percentage) as fixed, main effects, as well as all possible interactions between height, waist, sample, and sex. For all models, the continuous regressors involved in interactions (waist and height) were centred.

Although sample could be thought as a random factor (i.e. fitting linear mixed models instead), we treated it as a fixed effects categorical predictor in the models because there were only three levels (Bogota, Mexico City, Me’Phaa), and a minimum of five levels is recommended. To test the residual distribution, generalised linear models (GLM) were fitted, but in all cases, residuals were closer to a normal or gamma (inverse link) distribution, for each sample. Models here included were fitted using the *lm* function in R, version 3.6.1^65^.

The most parameterised initial model (Model 1) was then reduced, by excluding the main effects of hip, weight, fat percentage, visceral fat, BMI and muscle percentage (as these are phenotypic markers associated either with height or waist circumference), and keeping the main effects of age, as well as the main effects and all possible interactions between any combination of height, waist, sample, and sex, consistent with our predictions. This, still highly parameterised model (Model 2), was further reduced using the functions *dredge* (https://www.rdocumentation.org/packages/MuMIn/versions/1.43.6/topics/dredge) and *model.sel* (https://www.rdocumentation.org/packages/MuMIn/versions/1.43.6/topics/model.sel) from the package *MuMIn: Multi-Model Inference*^66^. The *dredge* function fitted a set of 334 models with combinations (subsets) of fixed effect terms from the second model, that were then compared using the function *model.sel* based on the Akaike Information Criterion (AICc) and Akaike weights, allowing us to select the best model (Model 3). This, best-supported model (i.e. the model with the lowest ΔAICc with a AICc higher than two units from the second most adequate model), is reported^67^.

Finally, we compared the three models selected model (Models 1, 2 and 3) using the *ICtab* function from the *bbmle* package^68^. Once a final model was selected, model diagnostics were performed (collinearity, residual distribution and linearity of residuals in each single term effect; see section 3.3 in the Supplementary Material).

Interactions in the final model were explored and via simple slopes analysis and Johnson-Neyman intervals^69, 70^, using the R package *interactions: Comprehensive, User-Friendly Toolkit for Probing Interactions*^71^. For this purpose, we implemented the functions *sim_slopes* (https://www.rdocumentation.org/packages/interactions/versions/1.1.1/topics/sim_slopes), *interact_plot* (https://www.rdocumentation.org/packages/interactions/versions/1.1.1/topics/interact_plot) and *johnson_neyman* (https://www.rdocumentation.org/packages/interactions/versions/1.1.1/topics/johnson_neyman).

## Results

All data and code used to perform these analyses are openly available from the Open Science Framework (OSF) project for this study (https://osf.io/5rzfs/).

### Descriptives

Descriptive statistics of age, waist circumference, hip, height, weight, fat percentage, visceral fat, BMI, muscle percentage and self-reported health and reported in Table 1.

**Table 1.**
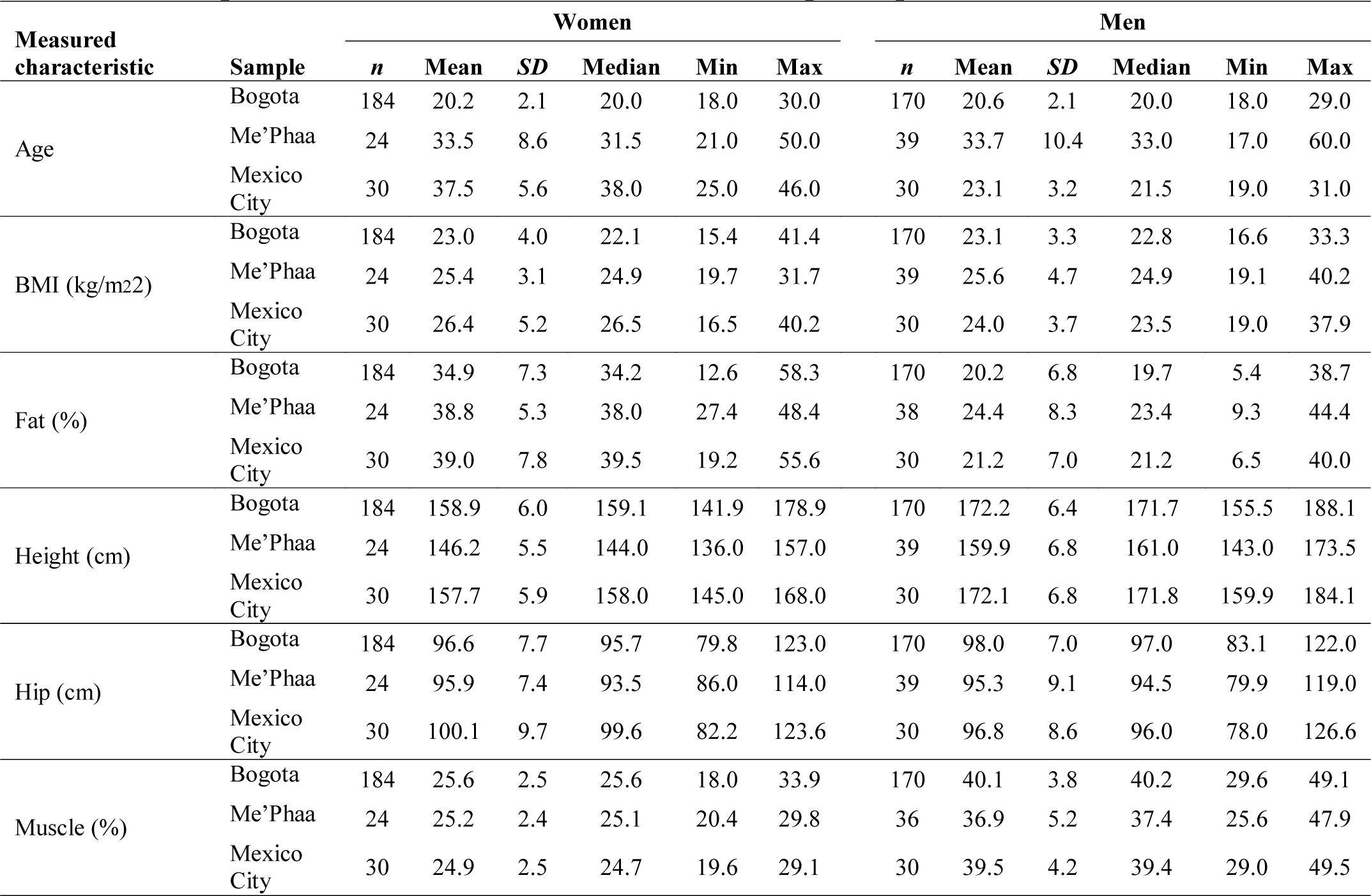
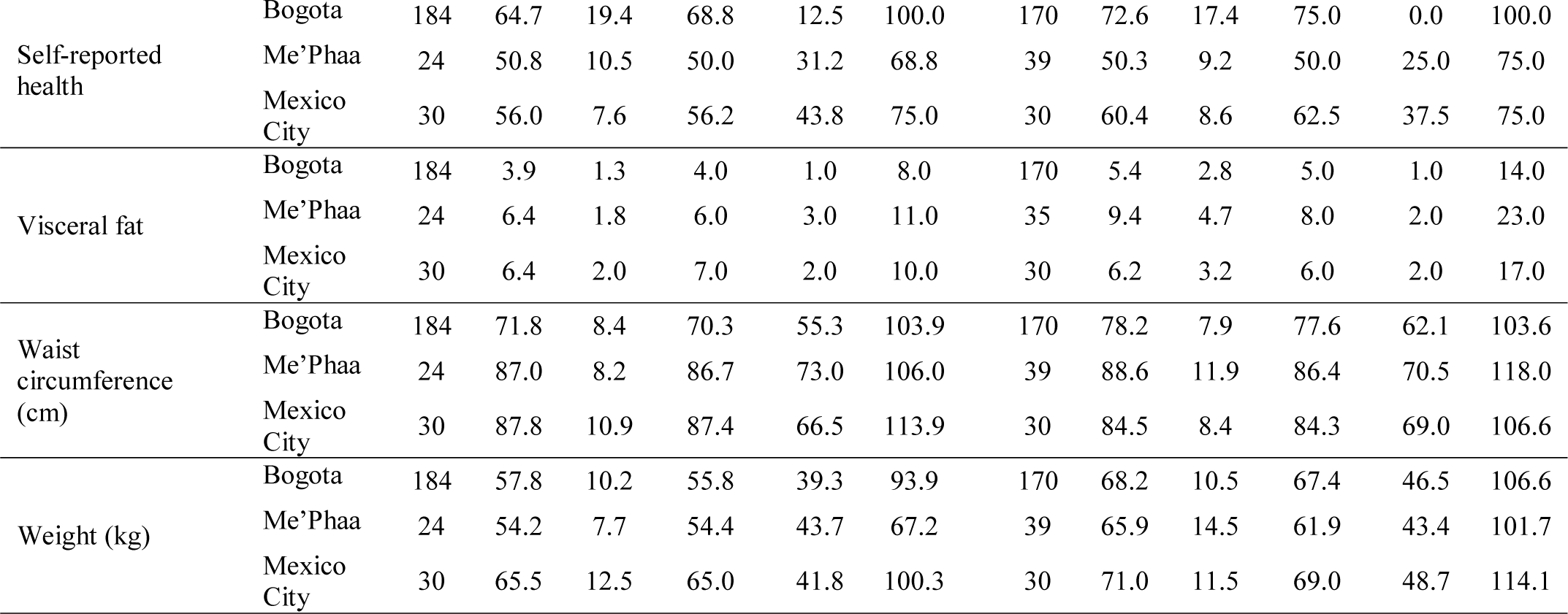
Descriptive statistics of measured variables of all participants.

The distribution of all measured variables is shown in Figure 1. Age, waist, height, visceral fat, and self-reported health strongly varied in both women (Fig 1a) and men (Fig 1b) between samples.

**Figure 1.**
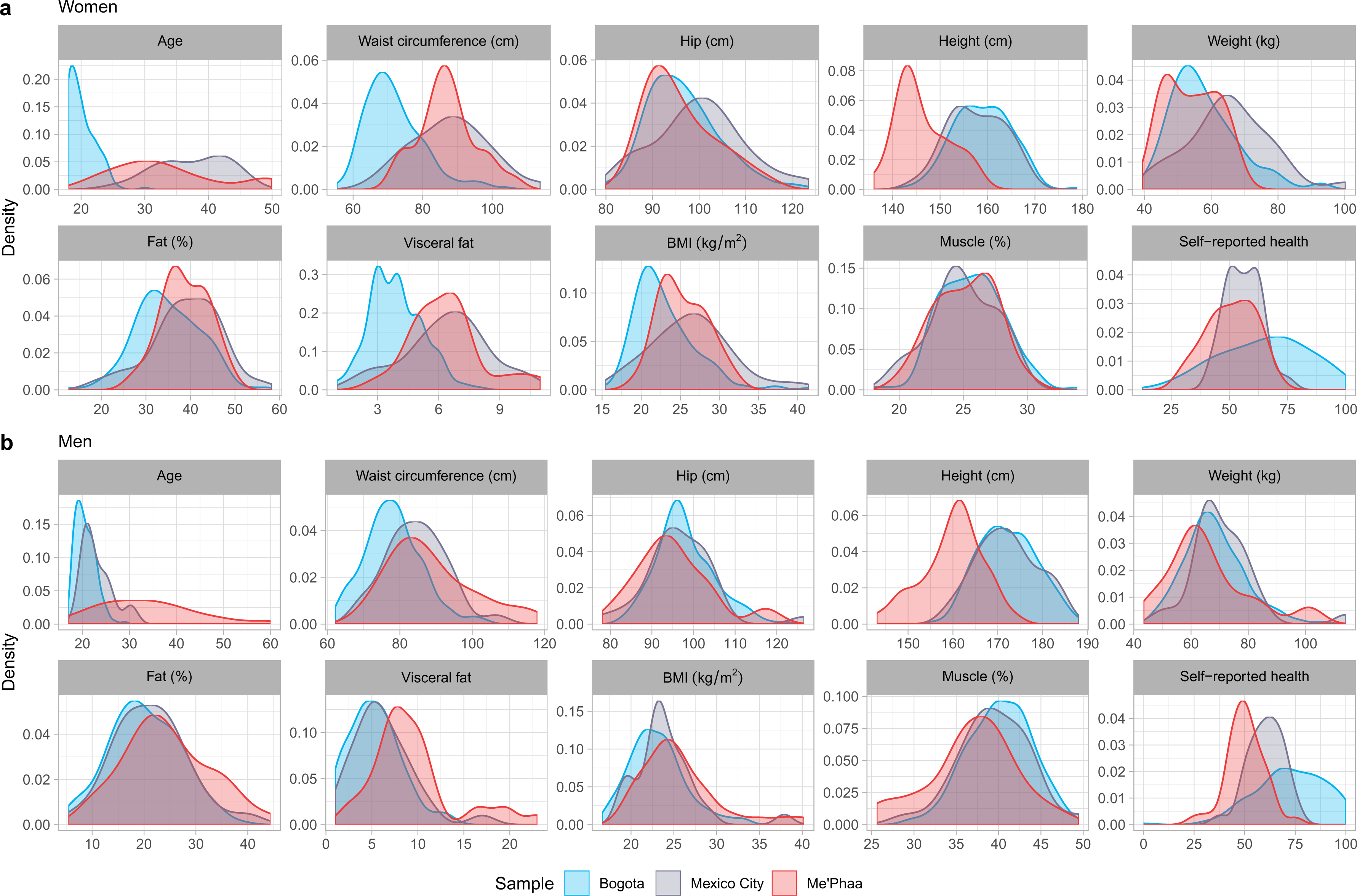
Distribution of all measured variables by sex and sample. (**a**) Women. (**b**) Men. For a comparison of sex differences in height, waist circumference and self-reported health in the three samples, see Supplementary Figure S1 online.

Age, waist circumference, height, fat percentage, visceral fat, BMI and muscle percentage, were significantly correlated with self-reported health (*r* > 0.10, in all cases) in both men and women (bivariate Pearson correlations between all measured variables for all participants combined are shown in the Supplementary Table S2, for women in Supplementary Table S3, and for men in Supplementary Table S4, online).

**Table 2.**
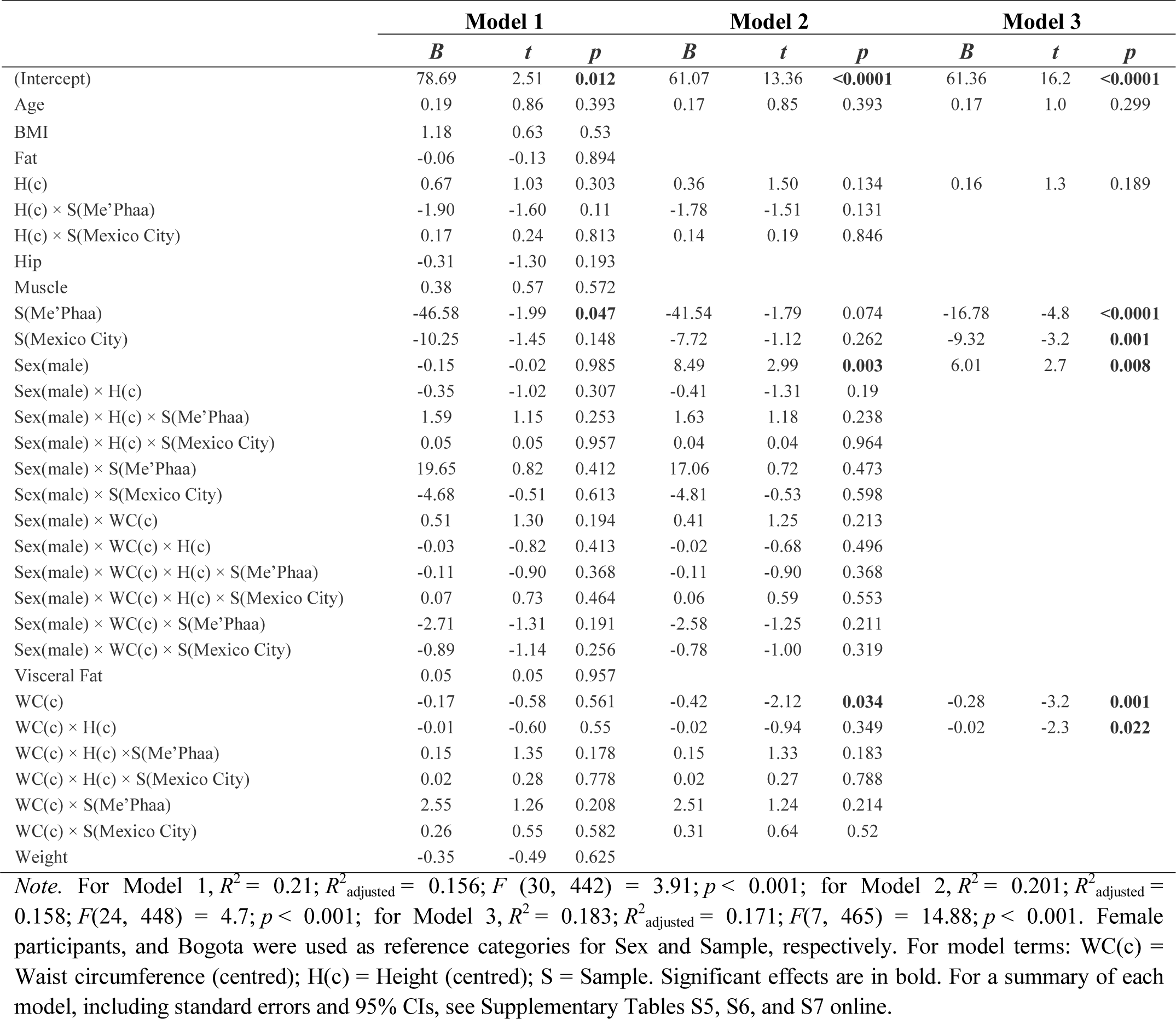
Results of separate LMs testing effects of independent variables on self-reported health.

|  | Model 1 |  |  | Model 2 |  |  | Model 3 |  |  |
| --- | --- | --- | --- | --- | --- | --- | --- | --- | --- |
|  | *B* | *t* | *p* | *B* | *t* | *p* | *B* | *t* | *p* |
| (Intercept) | 78.69 | 2.51 | **0.012** | 61.07 | 13.36 | **<0.0001** | 61.36 | 16.2 | **<0.0001** |
| Age | 0.19 | 0.86 | 0.393 | 0.17 | 0.85 | 0.393 | 0.17 | 1.0 | 0.299 |
| BMI | 1.18 | 0.63 | 0.53 |  |  |  |  |  |  |
| Fat | -0.06 | -0.13 | 0.894 |  |  |  |  |  |  |
| H(c) | 0.67 | 1.03 | 0.303 | 0.36 | 1.50 | 0.134 | 0.16 | 1.3 | 0.189 |
| H(c) × S(Me'Phaa) | -1.90 | -1.60 | 0.11 | -1.78 | -1.51 | 0.131 |  |  |  |
| H(c) × S(Mexico City) | 0.17 | 0.24 | 0.813 | 0.14 | 0.19 | 0.846 |  |  |  |
| Hip | -0.31 | -1.30 | 0.193 |  |  |  |  |  |  |
| Muscle | 0.38 | 0.57 | 0.572 |  |  |  |  |  |  |
| S(Me'Phaa) | -46.58 | -1.99 | **0.047** | -41.54 | -1.79 | 0.074 | -16.78 | -4.8 | **<0.0001** |
| S(Mexico City) | -10.25 | -1.45 | 0.148 | -7.72 | -1.12 | 0.262 | -9.32 | -3.2 | **0.001** |
| Sex(male) | -0.15 | -0.02 | 0.985 | 8.49 | 2.99 | **0.003** | 6.01 | 2.7 | **0.008** |
| Sex(male) × H(c) | -0.35 | -1.02 | 0.307 | -0.41 | -1.31 | 0.19 |  |  |  |
| Sex(male) × H(c) × S(Me'Phaa) | 1.59 | 1.15 | 0.253 | 1.63 | 1.18 | 0.238 |  |  |  |
| Sex(male) × H(c) × S(Mexico City) | 0.05 | 0.05 | 0.957 | 0.04 | 0.04 | 0.964 |  |  |  |
| Sex(male) × S(Me'Phaa) | 19.65 | 0.82 | 0.412 | 17.06 | 0.72 | 0.473 |  |  |  |
| Sex(male) × S(Mexico City) | -4.68 | -0.51 | 0.613 | -4.81 | -0.53 | 0.598 |  |  |  |
| Sex(male) × WC(c) | 0.51 | 1.30 | 0.194 | 0.41 | 1.25 | 0.213 |  |  |  |
| Sex(male) × WC(c) × H(c) | -0.03 | -0.82 | 0.413 | -0.02 | -0.68 | 0.496 |  |  |  |
| Sex(male) × WC(c) × H(c) × S(Me'Phaa) | -0.11 | -0.90 | 0.368 | -0.11 | -0.90 | 0.368 |  |  |  |
| Sex(male) × WC(c) × H(c) × S(Mexico City) | 0.07 | 0.73 | 0.464 | 0.06 | 0.59 | 0.553 |  |  |  |
| Sex(male) × WC(c) × S(Me'Phaa) | -2.71 | -1.31 | 0.191 | -2.58 | -1.25 | 0.211 |  |  |  |
| Sex(male) × WC(c) × S(Mexico City) | -0.89 | -1.14 | 0.256 | -0.78 | -1.00 | 0.319 |  |  |  |
| Visceral Fat | 0.05 | 0.05 | 0.957 |  |  |  |  |  |  |
| WC(c) | -0.17 | -0.58 | 0.561 | -0.42 | -2.12 | **0.034** | -0.28 | -3.2 | **0.001** |
| WC(c) × H(c) | -0.01 | -0.60 | 0.55 | -0.02 | -0.94 | 0.349 | -0.02 | -2.3 | **0.022** |
| WC(c) × H(c) × S(Me'Phaa) | 0.15 | 1.35 | 0.178 | 0.15 | 1.33 | 0.183 |  |  |  |
| WC(c) × H(c) × S(Mexico City) | 0.02 | 0.28 | 0.778 | 0.02 | 0.27 | 0.788 |  |  |  |
| WC(c) × S(Me'Phaa) | 2.55 | 1.26 | 0.208 | 2.51 | 1.24 | 0.214 |  |  |  |
| WC(c) × S(Mexico City) | 0.26 | 0.55 | 0.582 | 0.31 | 0.64 | 0.52 |  |  |  |
| Weight | -0.35 | -0.49 | 0.625 |  |  |  |  |  |  |

*Note.* For Model 1, $R^2 = 0.21$ ; $R^2_{\text{adjusted}} = 0.156$ ; $F(30, 442) = 3.91$ ; $p < 0.001$ ; for Model 2, $R^2 = 0.201$ ; $R^2_{\text{adjusted}} = 0.158$ ; $F(24, 448) = 4.7$ ; $p < 0.001$ ; for Model 3, $R^2 = 0.183$ ; $R^2_{\text{adjusted}} = 0.171$ ; $F(7, 465) = 14.88$ ; $p < 0.001$ . Female participants, and Bogota were used as reference categories for Sex and Sample, respectively. For model terms: WC(c) = Waist circumference (centred); H(c) = Height (centred); S = Sample. Significant effects are in bold. For a summary of each model, including standard errors and 95% CIs, see Supplementary Tables S5, S6, and S7 online.

**Table 3.** Performance criteria of the three selected models.

| Model | AICc | $\Delta\text{AICc}$ | $df$ | $w_i(\text{AICc})$ |
| --- | --- | --- | --- | --- |
| Model 3 | 3999 | 0 | 9 | 0.99999782 |
| Model 2 | 4025 | 26 | 26 | 0.00000215 |
| Model 1 | 4034 | 35 | 32 | 0.00000003 |

*Note.* Models are in descending order from the best to the worst fitting. $\Delta AIC$ is the change in AIC between each model and the best model. Akaike weights ( $w_i(AICc)$ ) are conditional probabilities for each model being the best model <sup>67</sup>.

**Table 4.** Results of the final LMM testing effects of independent variables on self-reported health

|  | *B* | *SE(B)* | 95% CI | *t* | *p* |
| --- | --- | --- | --- | --- | --- |
| (Intercept) | 61.36 | 3.78 | 53.94 — 68.782 | 16.25 | **<0.0001** |
| Age | 0.17 | 0.16 | -0.153 — 0.495 | 1.04 | 0.299 |
| H(c) | 0.16 | 0.12 | -0.08 — 0.401 | 1.31 | 0.189 |
| S(Mexico City) | -9.32 | 2.87 | -14.96 — -3.681 | -3.25 | **0.001** |
| S(Me'Phaa) | -16.78 | 3.53 | -23.722 — -9.83 | -4.75 | **<0.0001** |
| Sex(male) | 6.01 | 2.26 | 1.564 — 10.458 | 2.66 | **0.008** |
| WC(c) | -0.28 | 0.09 | -0.459 — -0.11 | -3.20 | **0.001** |
| WC(c) × H(c) | -0.02 | 0.01 | -0.036 — -0.003 | -2.30 | **0.022** |
| **Simple slope analysis for H(c) at different values of WC(c)** |  |  |  |  |  |
| WC(c) - 1 SD = -10.53 | 0.37 | 0.15 | 0.075 — 0.657 | 2.47 | **0.014** |
| WC(c) Mean = -0.12 | 0.16 | 0.12 | -0.077 — 0.403 | 1.33 | 0.183 |
| WC(c) + 1 SD = 10.28 | -0.04 | 0.15 | -0.341 — 0.261 | -0.26 | 0.795 |

*Note.* As waist reference, the centred values used are equivalent to -1 SD (67.43 cm), mean (78.01 cm), and +1 SD (88.6 cm). Women and Bogota were used as reference categories for Sex and Sample, respectively. For model terms: WC(c) = Waist circumference (centred); H(c) = Height (centred); S = Sample. Significant predictors are in bold.

### Models to predict self-reported health

To establish the relationship between height and self-reported health, we fitted three linear models (Table 2). For all models, the continuous regressors involved in interactions (waist and height) were mean-centred.

In the first model (Model 1), we included as predictors all measured variables as main effects, as well as all interactions between height, waist circumference, sample, and sex. The first model was initially reduced by excluding hip, weight, fat percentage, BMI and muscle percentage. We decided to include waist circumference instead of visceral fat or fat percentage for two reasons: first, because these three variables are strongly correlated in women and men (*r* > 0.79 in all cases; see Supplementary Tables S3 and S4 online, for women and men, respectively). And second, because unlike visceral fat or fat percentage, waist circumference can be directly perceived by others, and hence could have a direct effect on mate-choice; fat percentage and visceral fat, on the other hand, are likely perceived and assessed in social contexts through other variables, including relative waist size.

In the second model (Model 2), we therefore included age, height, sample, sex, waist circumference, and all possible interactions between combinations of height, waist circumference, sample, and sex. This second model was further reduced by the implementation of the functions *dredge* and *model.sel* from the package MuMIn^66^ (for details, see the Statistical analysis section in the Methods). These functions fitted and compared a total of 334 models with different combinations of fixed terms from Model 2; these compared models and their relative probability to be the best model are shown according to their relative Akaike weights (*w_i_*(AICc)) in Fig. 2.

**Figure 2.**
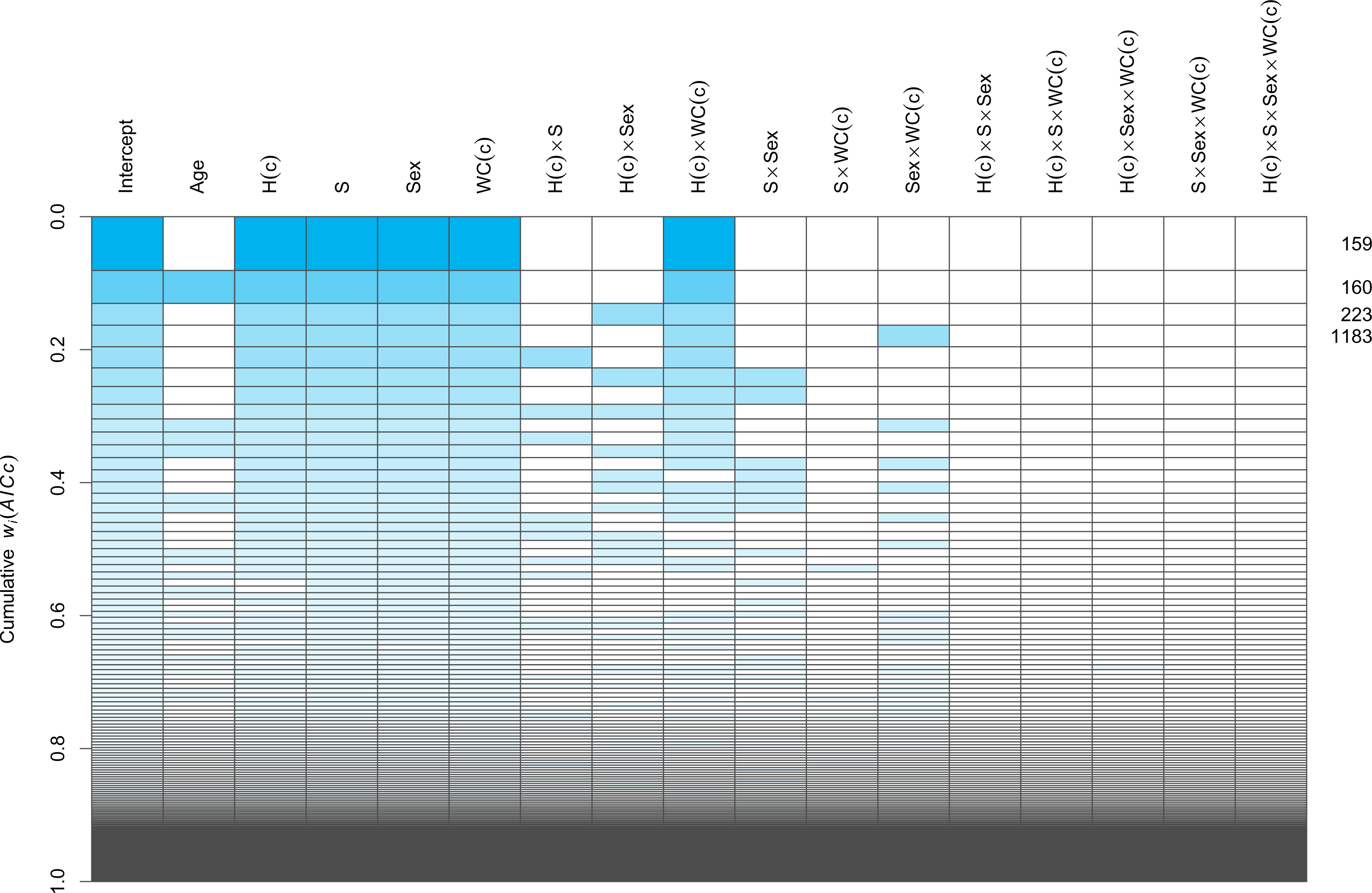
Model selection plot. Rows represent each of the 334 compared models. Cells coloured in blue represent included terms in each model, according to their Akaike weight (*wi*(*AICc*)), represented as the height of each row/model. Given the important age differences between samples, we selected the second-best model (labelled **160**), because it had the same structure as the best model (labelled **159**), but also included Age as a regressor. Furthermore, this second-best model had a Δ*AICc* of less than 2 units (≈ 0.98) compared to the best model. For model terms: WC(c) = Waist circumference (centred); H(c) = Height (centred); S = Sample.

This analysis revealed that the best model (labelled 159 in Fig. 2), included height (centred), sample, sex, waist circumference (centred) and the interaction between height (centred) and waist (centred). However, to account for the age differences between samples, we selected the second-best model (labelled 160 in Fig. 2), because it also included age as a regressor, and had a ΔAICc of less than 2 units (≈ 0.98) compared to the best model. This model, including age, was therefore selected as our final model (Model 3).

The three selected models were compared using the AICc, Akaike weights (*w_i_*(AICc)) and ΔAICc (Table 3). The analyses revealed that Model 3 is not only the most parsimonious of the three selected models, but has higher *R*^2^_adjusted_ and *F* values (Table 2), as well as a lower AIC and higher Akaike weight^67^ (Table 3) than the previous two models. In fact, Model 3 is close to 464,686 times more likely to be the best model compared to Model 2, and about 35,141,683 times compared to Model 1 (Model 2, was around 76 times more likely compared to Model 1).

Furthermore, for Model 3 (the final, minimum adequate model), Generalised Variance Inflation Factors (GVIF)^66^ revealed no concerning cases of collinearity for any of the predictor terms (GVIF ≤ 3, and a GVIF^1/(2×Df)^ ≤ 1.6 in all cases; for details, see Supplementary Table S8 online; residual distribution by sample and linearity in each single term factor are shown in Supplementary Fig. S2 online).

The final model (Model 3: Table 4; Fig. 3) showed a significant, negative main effect of waist circumference (*t* = −3.20, *p* = 0.001), and significant main effects of sex (men rated their health 6.01 points higher than women; *t* = 2.66, *p* = 0.008), and sample (Mexico City and Me’Phaa individuals rated their health 9.32 and 19.78 points lower than participants from Bogota; *t* = −3.25, *p* = 0.001 and; *t* = −4.75, *p* < 0.001, respectively).

**Figure 3.**
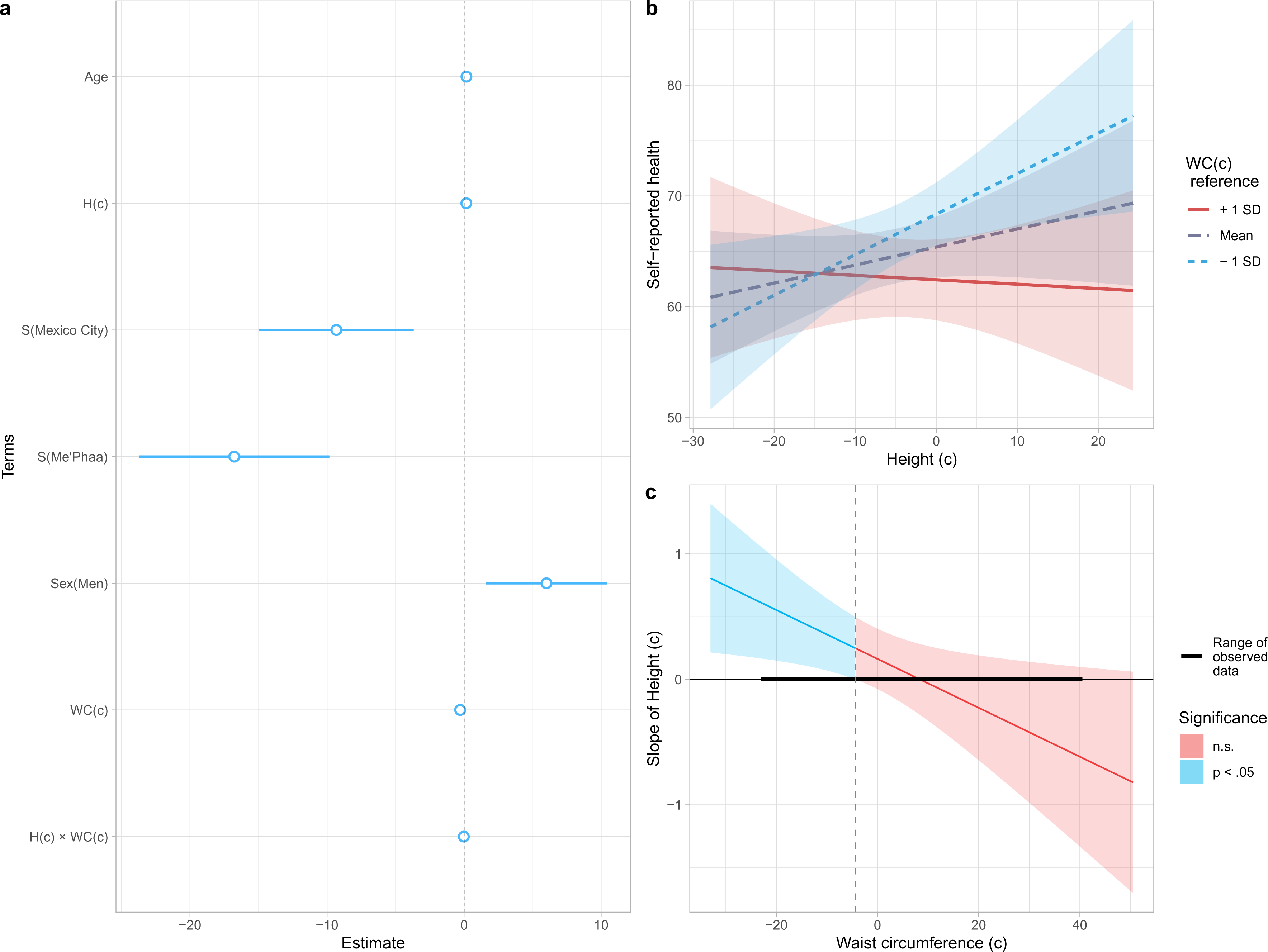
Model 3 estimates and interaction between Height and Waist. Values of Height and Waist were centred: for Height, mean ± SD = 163.83 ± 9.85; for Waist circumference, mean ± SD = 78.01 ± 10.59. (**a**) Estimates and 95% CI for each model term. For categorical predictors, women and Bogota were used as reference levels. For model terms, WC(c) = Waist circumference (centred); H(c) = Height (centred); S = Sample. (**b**) Interaction between Height and Waist. As waist reference, −1 SD (67.43 cm), mean (78.01 cm), and +1 SD (88.6 cm) values were used, showed on a blue to red colour scale. (**c**) Johnson-Neyman plot, showing for which values of Waist (centred), the slope of Height (centred) is significant as a predictor of Self-reported health; these slopes are predicted to be significant for centred Waists circumferences below −4.51 (73.51 cm), or above 68.73 (146.74 cm; not shown as it is a prediction for extreme values, beyond the ones found in any of our samples).

Moreover, a significant interaction between waist and height (Table 4; *t* = −2.30, *p* = 0.022) was revealed, indicating that the negative association of waist circumference with self-reported health was height-dependent (Fig. 3b); the best predicted self-reported health was for tall participants with small waists, and while the association between height and self-reported health is positive for people with small waist circumferences, it decreases for people with increasingly large waists. Furthermore, the Johnson-Neyman procedure^69, 70^ (Fig. 3c), indicated that height is only a significant, always positive, predictor of self-reported health for people with relatively small waist circumferences of less than 73.51 cm (centred: −4.51).

This interaction was replicated when fitting an alternative version of Model 3 (Model 3A), replacing waist circumference for visceral fat, by following the same method to select it (i.e. fitting an alternative Model 2, and repeating the same selection process; see section 4.3 in the Supplementary Material, online). Similar to Model 3, this alternative Final Model, also included an interaction between height and, in this case, visceral fat, in which height was found to be a positive significant predictor of self-reported health, only for people with low levels of visceral fat (see Table S9 and Fig. S3 in the Supplementary Material, online). Furthermore, in this model the interaction was more extreme than when using waist circumference, and height becomes a significant, negative predictor of self-reported health for people with high visceral fat (see Fig. S3c in the Supplementary Material, online).

## Discussion

The present study provides new insights into the relationship between height and health in men and women by studying three Latin-American populations, which included urban and indigenous populations with marked differences in access to basic needs and services like food and health.

Contrary to our initial hypothesis, height was not a significant predictor of self-perceived health but interacted with waist circumference. Most results in favour of a direct relationship between height itself and health were carried out in small modern populations and specific Western ethnic groups more than twenty years ago. New studies with non-traditional population groups have failed to verify the positive relationship between height and health, especially associated with cardiovascular and autoimmune diseases^72, 73^. For example, studies on Native Americans, Japanese, Indians and Pakistanis showed that shorter people had a lower prevalence of cardiovascular disease than the tallest people in each population^73^. These findings were similar in Sardinian inhabitants, a European population with the lowest physical stature recorded in Europe in recent years^72^.

Interestingly, our results suggest that there is a main negative effect of waist circumference on self-perceived health. This is congruent with a broad range of studies done in different human populations^74^. In fact, waist circumference has been proposed as one of the most important biomarker of metabolic syndrome that predicts health condition in terms of cardiovascular diseases^60^. Nevertheless, we found that this negative association was height-dependent in our studied samples. In other words, waist circumference predicted self-reported health differently for people of different heights: while being taller predicts better self-reported health for taller people with relatively small waists, being taller was found to be associated with poorer perceptions of their health in people with larger waist circumferences. Furthermore, while there is a cost of abdominal and visceral adiposity for tall people, there is no predicted cost for shorter persons. Interestingly, epidemiological studies have widely implemented an integration of both phenotypical components in the form of waist to height ratio (WtHR). In general, waist circumference has stronger negative impact on health of short individuals than for tall ones^75^, contrasting with our results. These differences might be due to WtHR has been mainly used to predict health in terms of metabolic and cardiovascular diseases (CVD), while our study used a general status of health, which could include more than metabolic and CVD. In addition, we use these phenotypic variables as continuous and independent predictors because the aim of our study argued that human height by itself would be an honest indicator of general health, which would not be able to be evaluate with WtHR as predictor. Therefore, our results argue the importance of considering a phenotypic independent integration of different human features that could be involved in health or physiological conditions, when a possible sexually selected trait is being evaluated as a signal of individual condition.

On the other hand, given that height is the human anatomical feature most sensitive to environmental and socioeconomic conditions^18, 38^, we expected stronger association between health and height for the indigenous population where the cost to produce and maintain this costly trait is greater than for inhabitants from urbanised areas. Nevertheless, we did not find inter-population differences in the magnitude of this relation. Urban populations reported better health than the indigenous population, and the shortest participants tended to be from the indigenous Me’Phaa sample. These results could, in fact, suggest different life-history strategies. Compared with modern Western societies, different life strategies could take place in harsh environments^76^, for instance, investing relatively less energy in growth and reallocating it towards reproduction^18^. In addition, a relative increase in the intensity or number of infectious diseases (including paediatric diseases in Me’Phaa) and higher tendency to early sexual maturity could negatively impact growth, resulting in a lower average height^77, 78^. These trends could be compensations between life-history components^35^. Finally, fast and prolonged growth imply high costs for the organism^1^. Rapid growth may influence mortality risk^79^ and growing for a longer time delays the onset of reproduction, increasing the risk of death and producing fewer offspring^1^. This perspective of life strategies allows us to understand the relationship between height, health and reproduction. This suggests the importance of addressing factors such as ethnicity, socioeconomic status, level of urbanisation in populations where there is great heterogeneity in access to food, health and pressure resources for pathogens, for instance, in Latin-American populations in which this relationship has barely been directly explored.

Although our results show that height and waist circumference are important predictors of self-perception of health, we did not evaluate any immunological mechanism that may underlie the self-perception responses of the participants. This limitation makes hard to directly evaluate human height as an honest signal of individual condition. In the present study, the questions done to evaluate general health (SF-36) is far to be a direct indicator of immune condition, since the participant’s perception responses could be influenced by components different than the individual’s ability to deal or resist to infectious pathogens, such as skeletal disorders, cancers, cardiovascular or metabolic abnormalities. Nevertheless, studies that have evaluated a more direct approximation of immune condition have led to controversial results. For instance, the implementation of antibody response to a hepatitis-B vaccine as a marker of immune condition has been positive associated with height in men (but not in women) up to a height of 185 cm, but an inverse relationship in taller men^80^. Furthermore, this relation was not found when different components of innate and adaptive immune system functioning were evaluated, such as lysozyme activity, neutrophil function, IgA and IgG^81^.

Finally, in relation with sex differences, women reported lower average health than men in all communities, which is concordant with reports and normative SF-36 data in other populations, especially in younger people^82, 83^. These results could consolidate the idea that height is a reliable signal of health in men^35^, while it could reflect reproductive success in women^84^ in terms of labour and birth, and to a lesser extent, function as an indicator of health^85^. It has been seen that taller women experience fewer problems during the labour process due to a lower risk of mismatch between foetal head size and size of the birth canal^85^. Nevertheless, this speculative idea warrants further studies on comparing health, reproductive success and female height.

It is important to consider that the mode of survey administration may be another limitation in our study, and it could have led to confounding effects. For example, it is possible that indigenous people have different understanding and thresholds about their general health perception, which we were unable to evaluate without previous validation of translated items, and it could have explained the lowest values of general health reported by indigenous people. Nevertheless, it could also reflect the real health conditions in Me’Phaa communities and not a misunderstanding of the survey. Other national indicators of health, such as morbidity and mortality by gastrointestinal and nasopharyngeal infectious diseases, have reported that Me’Phaa communities also present the poorest health in Mexico^52^, which is consistent with our results. In fact, items for the dimension of general health perception have the lowest standard deviation and coefficient of variation in the entire SF-36 survey, in both validated Spanish^55, 58^ and English versions^86^, which makes this dimension the most understandable one.

In addition, in order to consider obvious differences in language and perception of health, statistical models in this study assumed these inter-population variations *a priori*. The effects of the sample were considered in all performed LMs. We found that although samples differ considerably, the associations between height, waist circumference and self-perceptions of health were predicted to be in the same direction for all populations (i.e. not interacting with the sample).

Finally, we did not have any information regarding potential pregnancy history in women. This is important because each pregnancy can affect waist circumference, so future studies should collect and control or include this variable in all fitted models.

The present study contributes information which could be important in the framework of human sexual selection. If health and genetic quality cues play an important role in human mate-choice^87^, and height and waist interact to signal health, its evolutionary consequences, including cognitive and behavioural effects, should be addressed in future research. This could be done by studying the interaction between waist circumference and height, in relation to reproductive and/or mating success, as well as mate preferences and perceived attractiveness, in populations with both Westernised and non-Westernised lifestyles.

## Data availability

All data used for this article are openly available at the OSF^88^. Code to perform all analyses, data manipulation, tables and figures is available in PDF (‘Supplementary_Material.pdf’) and *R Markdown* (‘Supplementary_Material.Rmd’) formats, so that it can be fully reproduced and explored in depth^89^.

## Supporting information

Supplementary Material: Code and Analyses

## Author contributions

JDL, ORS, MV-A, EV and I.G-S. conceived and designed this study. JDL, ORS, AC-C, LM-S, and IG-S collected data. JDL and IG-S analysed all data. JDL, ORS, MV-A and IG-S wrote the first draft. All authors contributed to writing, approved the final version of the manuscript and gave approval for publication.

## Acknowledgments

We are grateful to Laura Rojas, Angie Ramos, Ángela Valderrama, Valentina West, Sergio Camelo, Laura Quintero, Paula Garzón, María Aguirre, Andrea Pastrana, Nicola Caro, Irene Olivella. Luisa Ramírez, Laura Guarín, y Henry Segura for their help in data collection in Colombia, and all our participants. We also grateful to Xuajin Me’Phaa, Margarita Mucino, Julio Gatica, and Diego Hernandez-Mucino for their help in the liaison with the Me’Phaa community, and for their help in data collection logistics. This project was funded in Colombia by Colciencias [grant number C145I004800000881-1 to JDL] and Universidad El Bosque, Vice-rectory of Research [grant number PCI.2017-9444]. Logistics and data collection in Mexico were supported by UNAM-PAPIIT [grant numbers IA209416, IA207019] and CONACYT Ciencia Básica [grant number 241744].

## Competing interests

The authors declare that they have no competing interests.

## Notes

#### Summary of Updates

Main changes: 1. We have modified the conceptual framework along the whole manuscript, indicating that Self-perception of health is a very indirect proxy of immunocompentence. Hence, the term "immunocompentence" was treated more carefully and only as a possible, limited mechanism of general health perception. 2. Now, instead of using Linear Mixed Models, we use Linear Models, and include Sample (Bogota, Mexico City, Me'Phaa) as a fixed factor in all models, instead of the both Country (Colombia, Mexico) and Population (urban, indigenous). 3. We have added more information regarding the model selection process, as well as information of each model. Importantly, the main findings remain and indeed are strengthened, and even replicated when using visceral fat instead of waist circumference as an abdominal adiposity measure.

https://osf.io/kgr5x/

## References

1. Sear, R. Height and reproductive success□ : is bigger always better? in Homo Novus: A Human Without Illusions (eds. Frey, U. J., Störmer, C. & Willführ, K. P.) vol. 44 127–143 (Springer Berlin Heidelberg, 2010).

2. Pawlowski, B., Dunbar, R. I. M. & Lipowicz, A. Tall men have more reproductive success. Nature 403, 156 (2000).

3. Stulp, G., Buunk, A. P., Pollet, T. V., Nettle, D. & Verhulst, S. Are Human Mating Preferences with Respect to Height Reflected in Actual Pairings? PLoS One 8, e54186 (2013).

4. Salska, I. et al. Conditional mate preferences: Factors influencing preferences for height. Pers. Individ. Dif. 44, 203–215 (2008).

5. Sear, R., Allal, N. & Mace, R. Height, marriage and reproductive success in Gambian women. Res. Econ. Anthropol. 23, 203–224 (2004).

6. Silventoinen, K., Lahelma, E. & Rahkonen, O. Social background, adult body-height and health. Int. J. Epidemiol. 28, 911–918 (1999).

7. Manning, J. T. Fluctuating asymmetry and body weight in men and women: Implications for sexual selection. Ethol. Sociobiol. 16, 145–153 (1995).

8. Pawlowski, B. & Jasienska, G. Women’s preferences for sexual dimorphism in height depend on menstrual cycle phase and expected duration of relationship. Biol. Psychol. 70, 38–43 (2005).

9. Melamed, T. Personality correlates of physical height. Pers. Individ. Dif. 13, 1349–1350 (1992).

10. Blaker, N. M. et al. The height leadership advantage in men and women: Testing evolutionary psychology predictions about the perceptions of tall leaders. Gr. Process. Intergr. Relations 16, 17–27 (2013).

11. Peck, M. N. & Lundberg, O. Short stature as an effect of economic and social conditions in childhood. Soc. Sci. Med. 41, 733–738 (1995).

12. Mueller, U. & Mazur, A. Evidence of unconstrained directional selection for male tallness. Behav. Ecol. Sociobiol. 50, 302–311 (2001).

13. Nettle, D. Height and reproductive success in a cohort of british men. Hum. Nat. 13, 473–491 (2002).

14. Nettle, D. Women’s height, reproductive success and the evolution of sexual dimorphism in modern humans. Proc. R. Soc. B Biol. Sci. 269, 1919–1923 (2002).

15. Pawlowski, B. Variable preferences for sexual dimorphism in height as a strategy for increasing the pool of potential partners in humans. Proc. R. Soc. B Biol. Sci. 270, 709–712 (2003).

16. Re, D. E. & Perrett, D. I. Concordant preferences for actual height and facial cues to height. Pers. Individ. Dif. 53, 901–906 (2012).

17. Stearns, S. C. Life history evolution: successes, limitations, and prospects. Naturwissenschaften 87, 476–486 (2000).

18. Walker, R. et al. Growth rates and life histories in twenty-two small-scale societies. Am. J. Hum. Biol. 18, 295–311 (2006).

19. Samaras, T. T. How height is related to our health and longevity: A review. Nutr. Health 21, 247– 261 (2012).

20. Wells, J. The Thrifty Phenotype Hypothesis: Thrifty Offspring or Thrifty Mother? J. Theor. Biol. 221, 143–161 (2003).

21. Hayflick, L. & Moorhead, P. S. The serial cultivation of human diploid cell strains. Exp. Cell Res. 25, 585–621 (1961).

22. Giovannelli, L. et al. Nutritional and lifestyle determinants of DNA oxidative damage□: a study in a Mediterranean population. Carcinogenesis 23, 1483–1489 (2002).

23. Perry, R. J., Farquharson, C. & Ahmed, S. F. The role of sex steroids in controlling pubertal growth. Clin. Endocrinol. (Oxf). 68, 4–15 (2008).

24. Ellison, P. T. On fertile ground: A natural history of human reproduction. (Harvard University Press, 2009).

25. Iravani, M., Lagerquist, M., Ohlsson, C. & Sävendahl, L. Regulation of bone growth via ligand-specific activation of estrogen receptor alpha. J. Endocrinol. 232, 403–410 (2017).

26. Eco-immunology: Evolutive Aspects and Future Perspectives. (Springer, 2014). doi:10.1007/978-94-017-8712-3.

27. Folstad, I. & Karter, A. J. Parasites, bright males, and the immunocompetence handicap. Am. Nat. 139, 603–622 (1992).

28. Ansar Ahmed, S., Karpuzoglu, E. & Khan, D. Effects of Sex Steroids on Innate and Adaptive Immunity. in Sex Hormones and Immunity to Infection (eds. Klein, S. L. & Roberts, C.) 19–51 (Springer, 2010). doi:10.1007/978-3-642-02155-8_2.

29. Roved, J., Westerdahl, H. & Hasselquist, D. Sex differences in immune responses: Hormonal effects, antagonistic selection, and evolutionary consequences. Horm. Behav. 88, 95–105 (2017).

30. Bernin, H. & Lotter, H. Sex bias in the outcome of human tropical infectious diseases: Influence of steroid hormones. J. Infect. Dis. 209, (2014).

31. Neyrolles, O. & Quintana-Murci, L. Sexual Inequality in Tuberculosis. PLoS Med. 6, e1000199 (2009).

32. Nhamoyebonde, S. & Leslie, A. Biological Differences Between the Sexes and Susceptibility to Tuberculosis. J. Infect. Dis. 209, S100–S106 (2014).

33. Sheldon, B. C. & Verhulst, S. Ecological immunology: Costly parasite defences and trade-offs in evolutionary ecology. Trends Ecol. Evol. 11, 317–321 (1996).

34. Paajanen, T. A., Oksala, N. K. J., Kuukasjärvi, P. & Karhunen, P. J. Short stature is associated with coronary heart disease: a systematic review of the literature and a meta-analysis. Eur. Heart J. 31, 1802–1809 (2010).

35. Stulp, G. & Barrett, L. Evolutionary perspectives on human height variation. Biol. Rev. 91, 206– 234 (2016).

36. Wormser, D. et al. Adult height and the risk of cause-specific death and vascular morbidity in 1 million people: individual participant meta-analysis. Int. J. Epidemiol. 41, 1419–1433 (2012).

37. Henrich, J., Heine, S. J. & Norenzayan, A. The weirdest people in the world? Behav. Brain Sci. 33, 61–83 (2010).

38. Walker, R. & Hamilton, M. J. Life□History Consequences of Density Dependence and the Evolution of Human Body Size. Curr. Anthropol. 49, 115–122 (2008).

39. Deaton, A. Height, health, and development. Proc. Natl. Acad. Sci. 104, 13232–13237 (2007).

40. Garcia, J. & Quintana-Domeque, C. The evolution of adult height in Europe: A brief note. Econ. Hum. Biol. 5, 340–349 (2007).

41. Lim, S. S. et al. Measuring the health-related Sustainable Development Goals in 188 countries: a 30 baseline analysis from the Global Burden of Disease Study 2015. Lancet 388, 1813–1850 (2016).

42. Silventoinen, K. Determinants of variation in adult body height. J. Biosoc. Sci. 35, 263–285 (2003).

43. Dowd, J. B., Zajacova, A. & Aiello, A. Early origins of health disparities: Burden of infection, health, and socioeconomic status in U.S. children. Soc. Sci. Med. 68, 699–707 (2009).

44. Kuzawa, C. W. & Bragg, J. M. Plasticity in Human Life History Strategy. Curr. Anthropol. 53, S369–S382 (2012).

45. Bentham, J. et al. A century of trends in adult human height. Elife 5, e13410 (2016).

46. Human Development Report Office. Human Development Indicators and Indices: 2018 Statistical Update. http://hdr.undp.org/sites/default/files/2018_human_development_statistical_update.pdf (2018).

47. Fullman, N. et al. Measuring performance on the Healthcare Access and Quality Index for 195 countries and territories and selected subnational locations: a systematic analysis from the Global Burden of Disease Study 2016. Lancet 391, 2236–2271 (2018).

48. Poverty and inequality. Colombia Reports (2018).

49. Indigenous Peoples, Poverty and Human Development in Latin America. (The World Bank, 2004). doi:10.1596/978-1-4039-9938-2.

50. Montenegro, R. A. & Stephens, C. Indigenous health in Latin America and the Caribbean. Lancet 367, 1859–1869 (2006).

51. Biggs, B., King, L., Basu, S. & Stuckler, D. Is wealthier always healthier? The impact of national income level, inequality, and poverty on public health in Latin America. Soc. Sci. Med. 71, 266– 273 (2010).

52. SINAIS. Sistema Nacional de Informacion en Salud. http://www.sinais.salud.gob.mx (2016).

53. Miramontes, O., DeSouza, O., Hernández, D. & Ceccon, E. Non-Lévy Mobility Patterns of Mexican Me’Phaa Peasants Searching for Fuel Wood. Hum. Ecol. 40, 167–174 (2012).

54. Hernández-Muciño, D. et al. La comunidad me’phaa construye su futuro: agroecología y restauración como herramientas de desarrollo rural sustentable. in Experiencias de colaboración transdisciplinaria para la sustentabilidad (eds. Merçon, J., Ayala-Orozco, B. & Rosell, J. A.) 66–79 (CopIt ArXives, 2018).

55. Durán-Arenas, L., Gallegos-Carrillo, K., Salinas-Escudero, G. & Martínez-Salgado, H. Towards a Mexican normative standard for measurement of the short format 36 health-related quality of life instrument. Salud Publica Mex. 46, 306–15 (2004).

56. Duncan, P. T. The Morpho-Syntax of Indefinite Pronouns in Iliatenco Me’phaa. (University of Kansas, 2013).

57. Ware, J. E. & Sherbourne, C. D. The MOS 36-item short-form health survey (SF-36). I. Conceptual framework and item selection. Med. Care 30, 473–83 (1992).

58. Lugo A, L. H., García E, H. I. & Gómez R, C. Confiabilidad del cuestionario de calidad de vida en salud SF-36 en Medellín, Colombia. Rev. Fac. Nac. Salud Publica 24, 37–50 (2006).

59. Czernichow, S., Kengne, A.-P., Stamatakis, E., Hamer, M. & Batty, G. D. Body mass index, waist circumference and waist-hip ratio: which is the better discriminator of cardiovascular disease mortality risk? Evidence from an individual-participant meta-analysis of 82 864 participants from nine cohort studies. Obes. Rev. 12, 680–687 (2011).

60. Després, J. P. & Lemieux, I. Abdominal obesity and metabolic syndrome. Nature 444, 881–887 (2006).

61. Huxley, R., Mendis, S., Zheleznyakov, E., Reddy, S. & Chan, J. Body mass index, waist circumference and waist:hip ratio as predictors of cardiovascular risk—a review of the literature. Eur. J. Clin. Nutr. 64, 16–22 (2010).

62. Knowles, K. M., et al. Waist Circumference, Body Mass Index, and Other Measures of Adiposity in Predicting Cardiovascular Disease Risk Factors among Peruvian Adults. Int. J. Hypertens. 2011, 1–10 (2011).

63. Alberti, K. G. M., Zimmet, P. & Shaw, J. The metabolic syndrome—a new worldwide definition. Lancet 366, 1059–1062 (2005).

64. Expert Panel on Detection Evaluation and Treatment of High Blood Cholesterol in Adults. Executive Summary of The Third Report of The National Cholesterol Education Program (NCEP) Expert Panel on Detection, Evaluation, And Treatment of High Blood Cholesterol In Adults (Adult Treatment Panel III). JAMA 285, 2486–2497 (2001).

65. R Core Team. R: A language and environment for statistical computing. (2019).

66. Fox, J. & Monette, G. Generalized Collinearity Diagnostics. J. Am. Stat. Assoc. 87, 178–183 (1992).

67. Wagenmakers, E.-J. & Farrell, S. AIC model selection using Akaike weights. Psychon. Bull. Rev. 11, 192–196 (2004).

68. Bolker, B. bbmle: Tools for General Maximum Likelihood Estimation. R package version 1.0.20. (2017).

69. Johnson, P. O. & Fay, L. C. The Johnson-Neyman technique, its theory and application. Psychometrika 15, 349–367 (1950).

70. Bauer, D. J. & Curran, P. J. Probing Interactions in Fixed and Multilevel Regression: Inferential and Graphical Techniques. Multivariate Behav. Res. 40, 373–400 (2005).

71. Long, J. A. interactions: Comprehensive, User-Friendly Toolkit for Probing Interactions. R package version 1.1.0. (2019). 33

72. Pes, G. M. et al. The association of adult height with the risk of cardiovascular disease and cancer in the population of Sardinia. PLoS One 13, e0190888 (2018).

73. Samaras, T. T., Elrick, H. & Storms, L. H. Is short height really a risk factor for coronary heart disease and stroke mortality? A review. Med. Sci. Monit. 10, RA63–76 (2004).

74. Lean, M., Han, T. & Seidell, J. Impairment of health and quality of life in people with large waist circumference. Lancet 351, 853–856 (1998).

75. Schneider, H. J., Klotsche, J., Silber, S., Stalla, G. K. & Wittchen, H.-U. Measuring Abdominal Obesity: Effects of Height on Distribution of Cardiometabolic Risk Factors Risk Using Waist Circumference and Waist-to-Height Ratio. Diabetes Care 34, e7–e7 (2011).

76. Perry, G. H. & Dominy, N. J. Evolution of the human pygmy phenotype. Trends Ecol. Evol. 24, 218–225 (2009).

77. Harvey, P. H. & Clutton-Brock, T. H. Life History Variation in Primates. Evolution (N. Y). 39, 559–581 (1985).

78. Promislow, D. E. L. & Harvey, P. H. Living fast and dying young: A comparative analysis of life-history variation among mammals. J. Zool. 220, 417–437 (1990).

79. Rollo, C. D. Growth negatively impacts the life span of mammals. Evol. Dev. 4, 55–61 (2002).

80. Krams, I. A. et al. Body height affects the strength of immune response in young men, but not young women. Sci. Rep. 4, 1–3 (2014).

81. Pawlowski, B., Nowak, J., Borkowska, B., Augustyniak, D. & Drulis-Kawa, Z. Body height and immune efficacy: testing body stature as a signal of biological quality. Proc. R. Soc. B Biol. Sci. 284, 20171372 (2017).

82. Hopman, W. M. et al. Canadian normative data for the SF-36 health survey. CMAJ 163, 265–71 (2000).

83. Watson, E. K., Firman, D. W., Baade, P. D. & Ring, I. Telephone administration of the SF-36 health survey: validation studies and population norms for adults in Queensland. Aust. N. Z. J. Public Health 20, 359–363 (1996).

84. Gluckman, P. D. & Hanson, M. A. Evolution, development and timing of puberty. Trends Endocrinol. Metab. 17, 7–12 (2006).

85. Wells, J. C. K., DeSilva, J. M. & Stock, J. T. The obstetric dilemma: An ancient game of Russian roulette, or a variable dilemma sensitive to ecology? Am. J. Phys. Anthropol. 149, 40–71 (2012).

86. Walters, S. J. & Brazier, J. E. What is the relationship between the minimally important difference and health state utility values? The case of the SF-6D. Health Qual. Life Outcomes 1, 4 (2003).

87. Roberts, S. C. & Little, A. C. Good genes, complementary genes and human mate preferences. Genetica 132, 309–321 (2008).

88. Leongómez, J. D., Sánchez, O. R., Vásquez-Amézquita, M., Valderrama, E. & González-Santoyo, I. Data from: Self-reported Health is Related to Body Height and Waist Circumference in Rural Indigenous and Urbanised Latin-American Populations. Open Science Framework (2019) doi:10.17605/OSF.IO/KGR5X.

89. Leongómez, J. D., Sánchez, O. R., Vásquez-Amézquita, M., Valderrama, E. & González-Santoyo, I. Code and analyses for Self-reported Health is Related to Body Height and Waist Circumference in Rural Indigenous and Urbanised Latin-American Populations. Open Science Framework (2019) doi:10.17605/OSF.IO/9WKMT.

