## Supplementary Material: Code and Analyses for "Self-reported Health is Related to Body Height and Waist Circumference in Rural Indigenous and Urbanised Latin-American Populations"

*08 January, 2020*

**Description**

This `R Markdown` document contains the code and analyses for *Self-reported Health is Related to Body Height and Waist Circumference in Rural Indigenous and Urbanised Latin-American Populations* (2020). Data available at https://doi.org/10.17605/OSF.IO/KGR5X.

**4 Session info (for reproducibility) 37**

**5 Supplementary Reference 38**

### 1 Preliminaries

#### 1.1 Load Packages

Used packages include `osfr` to download and open data files directly from OSF (using the `osf_retrieve_file` and `osf_download` functions). Currently this package must be installed from GitHub. See instructions here. All other packages used in this file (full list in the code below) can be directly installed from the Comprehensive R Archive Network (CRAN).

```
library(osfr)
library(tidyverse)
library(gridExtra)
library(ggpubr)
library(psych)
library(kableExtra)
library(ryouready)
library(car)
library(survival)
library(jtools)
library(Hmisc)
library(lattice)
library(Formula)
library(bbmle)
library(lemon)
library(data.table)
library(rstatix)
library(MuMIn)
library(qqplotr)
library(interactions)
library(lmSupport)
```

#### 1.2 Custom functions

##### 1.2.1 Correlation matrix (`corstarsl`)

This function creates a correlation matrix, and displays significance (function `corstarsl` from MYOWELT).

```
corstarsl <- function(x) {
  require(Hmisc)
  x <- as.matrix(x)
  R <- rcorr(x)$r
```

```
  p <- rcorr(x)$P

  ## define notions for significance levels; spacing is important.
  mystars <- ifelse(p < .001,
                    "***",
                    ifelse(p < .01,
                           "** ",
                           ifelse(p < .05,
                                  "* ",
                                  " ")))

  ## trunctuate the matrix that holds the correlations to two decimal
  R <- format(round(cbind(rep(-1.11, ncol(x)), R), 2))[, -1]

  ## build a new matrix that includes the correlations with their apropriate stars
  Rnew <- matrix(paste(R, mystars,
                       sep = ""),
                 ncol = ncol(x))
  diag(Rnew) <- paste(diag(R), " ",
                      sep = "")
  rownames(Rnew) <- colnames(x)
  colnames(Rnew) <- paste(colnames(x), "",
                          sep = "")

  ## remove upper triangle
  Rnew <- as.matrix(Rnew)
  Rnew[upper.tri(Rnew, diag = TRUE)] <- ""
  Rnew <- as.data.frame(Rnew)

  ## remove last column and return the matrix (which is now a data frame)
  Rnew <- cbind(Rnew[1:length(Rnew) - 1])
  return(Rnew)
}
```

##### 1.2.2 Function to bold significant effects from model summary tables (summaSig)

This function formats and bolds significant *p*-values from model tables (e.g. `summary(model)$coefficients` and `anova(model)`).

```
summaSig <- function(modTab, pcol) {
  modTab[, pcol] <- ifelse(modTab[, pcol] < 0.0001,
                           "\\textbf{<0.0001}",
    ifelse(modTab[, pcol] < 0.001,
           "\\textbf{<0.001}",
      ifelse(modTab[, pcol] < 0.05,
             paste0("\\textbf{", round(modTab[, pcol], 3), "}"),
        round(modTab[, pcol], 3)
      )
    )
  )
  return(modTab)
}
```

##### 1.2.3 Function to format p-values (pvalr)

This function takes p-values and formats them (function `pvalr` from rawr).

```
pvalr <- function(pvals, sig.limit = .001, digits = 3, html = FALSE) {

  roundr <- function(x, digits = 1) {
    res <- sprintf(paste0('%.', digits, 'f'), x)
    zzz <- paste0('0.', paste(rep('0', digits), collapse = ''))
    res[res == paste0('-', zzz)] <- zzz
    res
  }

  sapply(pvals, function(x, sig.limit) {
    if (x < sig.limit)
      if (html)
        return(sprintf('&lt; %s', format(sig.limit))) else
          return(sprintf('< %s', format(sig.limit)))
    if (x > .1)
      return(roundr(x, digits = 2)) else
        return(roundr(x, digits = digits))
  }, sig.limit = sig.limit)
}
```

###### 1.2.4 Function format model terms from model summary tables (`summTerms`)

This function replaces term names from model tables (e.g.`summary(model)$coefficients`), and formats them into a shorter version.

```
summTerms <- function(summTab) {
  row.names(summTab) <- str_replace(row.names(summTab),
                                    "SexMen",
                                    "Sex(men)")
  row.names(summTab) <- str_replace(row.names(summTab),
                                    "Waist_c",
                                    "WC(c)")
  row.names(summTab) <- str_replace(row.names(summTab),
                                    "Height_c",
                                    "H(c)")
  row.names(summTab) <- str_replace(row.names(summTab),
                                    "SampleMexico City",
                                    "S(Mexico City)")
  row.names(summTab) <- str_replace(row.names(summTab),
                                    "SampleMe'Phaa",
                                    "S(Me'Phaa)")
  row.names(summTab) <- str_replace(row.names(summTab),
                                    "VisceralFat",
                                    "Visceral Fat")
  # This next line is repeated to make sure all interactions are dysplayed with
  # "×", instead of ":".
  row.names(summTab) <- str_replace(row.names(summTab),
                                    ":",
                                    " $\\\\times$ ")
  row.names(summTab) <- str_replace(row.names(summTab),
                                    ":",
                                    " $\\\\times$ ")
  row.names(summTab) <- str_replace(row.names(summTab),
                                    ":",
                                    " $\\\\times$ ")
  row.names(summTab) <- str_replace(row.names(summTab),
                                    ":",
```

```
                                         " $\\\\times$ ")
  return(summTab)
}
```

#### 1.3 Load and organise data from Colombia

##### 1.3.1 Anthropometric data

We collected 8 measures from each participant:

- Waist (circumference, in cm)
- Hip (circumference, in cm)
- Height (cm)
- Weight (kg)
- Fat (%)
- Visceral fat (score)
- BMI (kg/m$^2$)
- Muscle (%)

```
f1 <- osf_retrieve_file("q46sy") %>% osf_download(overwrite = TRUE)
mm <- read.csv(f1$local_path, sep = ",", dec = ".")
```

Because each anthropometric characteristic was measured 3 times, the intraclass correlation between measurements was assessed.

```
ICCwai <- ICC(mm[, 3:5]) # waist
ICChip <- ICC(mm[, 6:8]) # hip
ICChei <- ICC(mm[, 9:11]) # height
ICCwei <- ICC(mm[, 12:14]) # weight
ICCfat <- ICC(mm[, 15:17]) # fat percentage
ICCvisfat <- ICC(mm[, 18:20]) # visceral fat
ICCbmi <- ICC(mm[, 21:23]) # BMI
ICCmus <- ICC(mm[, 24:26]) # muscle percentage
```

###### 1.3.1.1 Table S1. Table displaying Intraclass Correlation for each measured characteristic

```
# paste ICC results
ICCtab <- rbind(ICCwai$results[1, ],
                ICChip$results[1, ],
                ICChei$results[1, ],
                ICCwei$results[1, ],
                ICCfat$results[1, ],
                ICCvisfat$results[1, ],
                ICCbmi$results[1, ],
                ICCmus$results[1, ])
# round numeric columns to 3 decimal places
ICCtab[, c(2, 3, 7, 8)] <- format(round(ICCtab[, c(2, 3, 7, 8)], 3),
                                  nsmall = 3)
# change rownames to column named "Anthropometric characteristic"
ICCtab <- ICCtab %>% rownames_to_column("Anthropometric characteristics")
# specify measured anthropometric characteristic
ICCtab$`Anthropometric characteristics` <- c("Waist circumference (cm)",
                                             "Hip (cm)",
                                             "Height (cm)",
                                             "Weight (kg)",
                                             "Fat (\\%)",
                                             "Visceral fat",
                                             "BMI (kg/m$^2$)",
```

```
                                                   "Muscle (\\%)")
# Clean table pasting CIs into one new column
ICCtab$ci <- paste0(ICCtab$`lower bound`, " - ", ICCtab$`upper bound`)
# replace p values for a more readable format
ICCtab <- summaSig(ICCtab, 7)
# select and reorder relevant columns
ICCtab <- ICCtab[, c(1, 3, 10, 4, 7)]

# print table with formatting
TabS1 <- kable(
  ICCtab,
  format = "latex",
  booktabs = TRUE,
  col.names = c("Anthropometric measure", "ICC", "95\\% CI", "$F$", "$p$"),
  caption = "\\textbf{Table S1.}
  Intraclass correlation of anthropometric characteristics measurements",
  align = c("l", "c", "c", "c", "c"),
  escape = FALSE) %>%
  kable_styling(latex_options = "HOLD_position") %>%
  footnote(
    general =
      "ICC values are ICC1
    (see \\\\href{https://tinyurl.com/yxjkdd44}{ICC function documentation}),
    which is a measure of absolute agreement. In all cases, $df$ were  353 and
    708. Significant results are in bold.",
    threeparttable = TRUE,
    escape = FALSE)
TabS1
```

**Table S1.** Intraclass correlation of anthropometric characteristics measurements

| Anthropometric measure | ICC | 95% CI | $F$ | $p$ |
|---|---|---|---|---|
| Waist circumference (cm) | 0.999 | 0.998 - 0.999 | 2339.214 | **<0.0001** |
| Hip (cm) | 0.998 | 0.998 - 0.999 | 1949.311 | **<0.0001** |
| Height (cm) | 0.999 | 0.999 - 0.999 | 4137.258 | **<0.0001** |
| Weight (kg) | 1.000 | 1.000 - 1.000 | 18901.869 | **<0.0001** |
| Fat (%) | 0.999 | 0.999 - 0.999 | 2553.452 | **<0.0001** |
| Visceral fat | 0.995 | 0.994 - 0.996 | 568.504 | **<0.0001** |
| BMI (kg/m$^2$) | 0.999 | 0.999 - 0.999 | 4157.237 | **<0.0001** |
| Muscle (%) | 0.999 | 0.998 - 0.999 | 2174.959 | **<0.0001** |

*Note:*
ICC values are ICC1 (see ICC function documentation), which is a measure of absolute agreement. In all cases, *df* were 353 and 708. Significant results are in bold.

Given the strong intraclass correlation between the three measurements of each anthropometric characteristic, we calculated the mean between those measurements.

```
mm$Waist <- rowMeans(mm[, 3:5])
mm$Hip <- rowMeans(mm[, 6:8])
mm$Height <- rowMeans(mm[, 9:11])
mm$Weight <- rowMeans(mm[, 12:14])
mm$Fat <- rowMeans(mm[, 15:17])
mm$VisceralFat <- rowMeans(mm[, 18:20])
mm$BMI <- rowMeans(mm[, 21:23])
```

```
mm$Muscle <- rowMeans(mm[, 24:26])
```

##### 1.3.2 Self-reported health data.

This data were obtained using a Spanish language validated translation[1] of the SF-36 questionnaire (https://www.rand.org/health-care/surveys_tools/mos/36-item-short-form.html). The translated version was validated in Colombia.

```
f2 <- osf_retrieve_file("p5cyf") %>% osf_download(overwrite = TRUE)
sf36 <- read.csv(f2$local_path, sep = ",", dec = ".")
```

The SF-36 produces 8 factors, calculated by averaging the recoded scores of individual items:

- Physical functioning (items 3 to 12): `PhysFunc`
- Role limitations due to physical health (items 13 to 16): `PhysLim`
- Role limitations due to emotional problems (items 17 to 19): `EmoLim`
- Energy/fatigue (items 23, 27, 29 and 31): `EnerFati`
- Emotional well-being (items 24, 25, 26, 28 and 30): `EmoWB`
- Social functioning (items 20 and 32): `SocFunc`
- Pain (items 21 and 22): `Pain`
- General health (items 1, 33, 34, 35 and 36): `Health`

To calculate this, all items were recoded following the instructions on how to score SF-36 (please see https://www.rand.org/health-care/surveys_tools/mos/36-item-short-form/scoring.html).

```
# New dataframe for recoded scores (and excluding ID)
sf36Re <- sf36[2:37]

# List of colnames
sf36Items <- colnames(sf36Re)

# Samples of items with same recoding
Reco1 <- sf36Items[c(1, 2, 20, 22, 34, 36)]
Reco2 <- sf36Items[c(3:12)]
Reco3 <- sf36Items[c(13:19)]
Reco4 <- sf36Items[c(21, 23, 26, 27, 30)]
Reco5 <- sf36Items[c(24, 25, 28, 29, 31)]
Reco6 <- sf36Items[c(32, 33, 35)]

# Recoding items by Sample
sf36Re <- recode2(sf36Re,
  vars = Reco1,
  recodes = "1 = 100;2 = 75;3 = 50;4 = 25;5 = 0")
sf36Re <- recode2(sf36Re,
  vars = Reco2,
  recodes = "1 = 0;2 = 50;3 = 100")
sf36Re <- recode2(sf36Re,
  vars = Reco3,
  recodes = "1 = 0;2 = 100")
sf36Re <- recode2(sf36Re,
  vars = Reco4,
  recodes = "1 = 100;2 = 80;3 = 60;4 = 40;5 = 20;6 = 0")
sf36Re <- recode2(sf36Re,
  vars = Reco5,
  recodes = "1 = 0;2 = 20;3 = 40;4 = 60;5 = 80;6 = 100")
sf36Re <- recode2(sf36Re,
  vars = Reco6,
  recodes = "1 = 0;2 = 25;3 = 50;4 = 75;5 = 100")
```

For the final factor calculation, we averaged recoded items (for detailed instructions, see https://www.rand.org/health-care/surveys_tools/mos/36-item-short-form/scoring.html). To make this data compatible with the Mexican database, and because item 35 cannot be answered by the Mexican Indigenous population, this item was excluded, and the `Health` factor was calculated averaging items 1, 33, 34, and 36 only.

```
sf36$SF.PhysFunc <- rowMeans(sf36Re[, 3:12])
sf36$SF.PhysLim <- rowMeans(sf36Re[, 13:16])
sf36$SF.EmoLim <- rowMeans(sf36Re[, 17:19])
sf36$SF.EnerFati <- rowMeans(sf36Re[, c(23, 27, 29, 31)])
sf36$SF.EmoWB <- rowMeans(sf36Re[, c(24:26, 28, 30)])
sf36$SF.SocFunc <- rowMeans(sf36Re[, c(20, 32)])
sf36$SF.Pain <- rowMeans(sf36Re[, 21:22])
sf36$SF.Health <- rowMeans(sf36Re[, c(1, 33, 34, 36)])
```

##### 1.4 Create final dataframe for the Colombian population

Combine the two data-frames into one, final data-frame (`dat`) containing only the relevant columns (i.e. the final factor scores for the SF-36 `Health` factor, and anthropometric means).

```
col <- merge(mm[, c(1, 2, 27:34)], sf36[, c(1, 38:45)], by = "ID")
```

The `Sex`, `Country` and `Population` columns were added. The `Sex` was based on the first letter of the ID of each participant (F = Female, M = Male). Finally, columns were reordered to match those of the Mexican database.

```
col$Sex <- NA
for (i in 1:length(col$Sex)) {
  col$Sex[i] <- ifelse(grepl("F", col$ID[i]), "Women", "Men")
}
col$Country <- "Colombia"
col$Population <- "Urban"
colS <- col[, c(1, 20:21, 19, 2:10, 18)]
```

##### 1.5 Load and organise data from Mexico

```
f3 <- osf_retrieve_file("kr8m9") %>% osf_download(overwrite = TRUE)
mex <- read.csv(f3$local_path, sep = ",", dec = ".")
```

Anthropometric data from Mexico had already been averaged from the three anthropometric measurements. We then calculated the `Health` factor from the SF-36 questionnaire, using the same system explained for the Colombian sample. (for detailed instructions on scoring of the SF-36 factors, see https://www.rand.org/health-care/surveys_tools/mos/36-item-short-form/scoring.html)

```
mex <- recode2(mex,
  vars = c("SF1", "SF34", "SF36"),
  recodes = "1 = 100;2 = 75;3 = 50;4 = 25;5 = 0")
mex <- recode2(mex,
  vars = "SF33",
  recodes = "1 = 0;2 = 25;3 = 50;4 = 75;5 = 100")
levels(mex$Sex) <- c("Women", "Men")
mex$SF.Health <- rowMeans(mex[, c(14:17)])
mexS <- mex[, c(1:13, 19)]
```

##### 1.6 Create final dataframe combining data from Colombian and Mexican samples

```
dat <- rbind(colS, mexS)
dat$Age <- as.numeric(dat$Age)
datcols <- c("Country", "Population", "Sex")
dat[datcols] <- lapply(dat[datcols], factor)
```

```
cols <- c("Country", "Population")
dat$Sample <- apply(dat[, cols], 1, paste, collapse = "-")
dat$Sample <- as.factor(dat$Sample)
colnames(dat)[14] <- "Health"
levels(dat$Sample) <- c("Bogota", "Me'Phaa", "Mexico City")
dat$Sample <- factor(dat$Sample, levels(dat$Sample)[c(1, 3, 2)])
dat$Sex <- factor(dat$Sex, levels(dat$Sex)[c(2, 1)])
write.csv(dat, file = "Full_data.csv", row.names = FALSE)
```

Data-frame structure

```
str(dat)
```

```
## 'data.frame':    477 obs. of  15 variables:
##  $ ID         : Factor w/ 477 levels "F001","F003",..: 1 2 3 4 5 6 7 8 9 10 ...
##  $ Country    : Factor w/ 2 levels "Colombia","Mexico": 1 1 1 1 1 1 1 1 1 1 ...
##  $ Population : Factor w/ 2 levels "Indigenous","Urban": 2 2 2 2 2 2 2 2 2 2 ...
##  $ Sex        : Factor w/ 2 levels "Women","Men": 1 1 1 1 1 1 1 1 1 1 ...
##  $ Age        : num  23 24 19 19 18 18 21 22 19 18 ...
##  $ Waist      : num  67.3 97.5 81.1 70.3 66.7 ...
##  $ Hip        : num  90.4 107.5 106.1 96.1 91.5 ...
##  $ Height     : num  158 165 165 161 162 ...
##  $ Weight     : num  48.8 71.4 73.9 56.2 54.8 ...
##  $ Fat        : num  30 42.6 43.5 34.2 32.4 ...
##  $ VisceralFat: num  3 5 5 3.67 3 ...
##  $ BMI        : num  19.7 26.2 27.1 21.7 20.9 ...
##  $ Muscle     : num  24.7 26.4 23.7 25.7 26.2 ...
##  $ Health     : num  75 50 43.8 50 68.8 ...
##  $ Sample     : Factor w/ 3 levels "Bogota","Mexico City",..: 1 1 1 1 1 1 1 1 1 1 ...
```

### 2 Descriptives

#### 2.1 Descriptives by Population, Sex and Country.

##### 2.1.1 Table 1. All participants

```
descColNames <- c(
    "Measured characteristic",
    "Sample",
    "$n$",
    "Mean",
    "$SD$",
    "Median",
    "Min",
    "Max")
descVarNames <- c(
  "Age",
  "Waist circumference (cm)",
  "Hip (cm)",
  "Height (cm)",
  "Weight (kg)",
  "Fat (\\%)",
  "Visceral fat",
  "BMI (kg/m$^2$)",
  "Muscle (\\%)",
  "Self-reported health")
```

```
# Subset of women participants
datF <- subset(dat, dat$Sex == "Women")
# Descriptives by Country and Population
descF <- describeBy(datF[5:14], datF$Sample, mat = TRUE, digits = 1)
# change rownames to column named "Measured characteristic"
descF <- descF[, c(2, 4:7, 10:11)] %>% rownames_to_column("Measured characteristic")
# specify measured anthropometric characteristic
varnames <- descVarNames
descF$`Measured characteristic` <- rep(varnames, each = 3)

# Subset of men participants
datM <- subset(dat, dat$Sex == "Men")
# Descriptives by Country and Population
descM <- describeBy(datM[5:14], datM$Sample, mat = TRUE, digits = 1)
# change rownames to column named "Measured characteristic"
descM <- descM[, c(2, 4:7, 10:11)] %>% rownames_to_column("Measured characteristic")
# specify measured anthropometric characteristic
varnames <- descVarNames
descM$`Measured characteristic` <- rep(varnames, each = 3)

# Merge Tables S2 and S3 by measured characteristic and sample
tab1 <- merge(descF, descM, by = c("Measured characteristic", "group1"), all = TRUE)

# Final fromated table
Tab1 <- kable(
  tab1,
  digits = 2,
  booktabs = TRUE,
  align = c("l", "l", rep("c", 12)),
  caption = "\\textbf{Table 1.}
  Descriptive statistics of measured variables for all participants",
  col.names = c("Measured characteristic", "Sample",
              rep(descColNames[3:8], 2)),
  escape = FALSE) %>%
  add_header_above(c(" " = 2,
                     "Women" = 6,
                     "Men" = 6)) %>%
  kable_styling(latex_options = c("scale_down", "HOLD_position"))  %>%
  collapse_rows(columns = 1, valign = "middle")
Tab1
```

**Table 1.** Descriptive statistics of measured variables for all participants

| | | Women | | | | | | Men | | | | | |
|---|---|---|---|---|---|---|---|---|---|---|---|---|---|
| Measured characteristic | Sample | *n* | Mean | *SD* | Median | Min | Max | *n* | Mean | *SD* | Median | Min | Max |
| Age | Bogota | 184 | 20.2 | 2.1 | 20.0 | 18.0 | 30.0 | 170 | 20.6 | 2.1 | 20.0 | 18.0 | 29.0 |
| | Me'Phaa | 24 | 33.5 | 8.6 | 31.5 | 21.0 | 50.0 | 39 | 33.7 | 10.4 | 33.0 | 17.0 | 60.0 |
| | Mexico City | 30 | 37.5 | 5.6 | 38.0 | 25.0 | 46.0 | 30 | 23.1 | 3.2 | 21.5 | 19.0 | 31.0 |
| BMI (kg/m$^2$) | Bogota | 184 | 23.0 | 4.0 | 22.1 | 15.4 | 41.4 | 170 | 23.1 | 3.3 | 22.8 | 16.6 | 33.3 |
| | Me'Phaa | 24 | 25.4 | 3.1 | 24.9 | 19.7 | 31.7 | 39 | 25.6 | 4.7 | 24.9 | 19.1 | 40.2 |
| | Mexico City | 30 | 26.4 | 5.2 | 26.5 | 16.5 | 40.2 | 30 | 24.0 | 3.7 | 23.5 | 19.0 | 37.9 |
| Fat (%) | Bogota | 184 | 34.9 | 7.3 | 34.2 | 12.6 | 58.3 | 170 | 20.2 | 6.8 | 19.7 | 5.4 | 38.7 |
| | Me'Phaa | 24 | 38.8 | 5.3 | 38.0 | 27.4 | 48.4 | 38 | 24.4 | 8.3 | 23.4 | 9.3 | 44.4 |
| | Mexico City | 30 | 39.0 | 7.8 | 39.5 | 19.2 | 55.6 | 30 | 21.2 | 7.0 | 21.2 | 6.5 | 40.0 |
| Height (cm) | Bogota | 184 | 158.9 | 6.0 | 159.1 | 141.9 | 178.9 | 170 | 172.2 | 6.4 | 171.7 | 155.5 | 188.1 |
| | Me'Phaa | 24 | 146.2 | 5.5 | 144.0 | 136.0 | 157.0 | 39 | 159.9 | 6.8 | 161.0 | 143.0 | 173.5 |
| | Mexico City | 30 | 157.7 | 5.9 | 158.0 | 145.0 | 168.0 | 30 | 172.1 | 6.8 | 171.8 | 159.9 | 184.1 |
| Hip (cm) | Bogota | 184 | 96.6 | 7.7 | 95.7 | 79.8 | 123.0 | 170 | 98.0 | 7.0 | 97.0 | 83.1 | 122.0 |
| | Me'Phaa | 24 | 95.9 | 7.4 | 93.5 | 86.0 | 114.0 | 39 | 95.3 | 9.1 | 94.5 | 79.9 | 119.0 |
| | Mexico City | 30 | 100.1 | 9.7 | 99.6 | 82.2 | 123.6 | 30 | 96.8 | 8.6 | 96.0 | 78.0 | 126.6 |
| Muscle (%) | Bogota | 184 | 25.6 | 2.5 | 25.6 | 18.0 | 33.9 | 170 | 40.1 | 3.8 | 40.2 | 29.6 | 49.1 |
| | Me'Phaa | 24 | 25.2 | 2.4 | 25.1 | 20.4 | 29.8 | 36 | 36.9 | 5.2 | 37.4 | 25.6 | 47.9 |
| | Mexico City | 30 | 24.9 | 2.5 | 24.7 | 19.6 | 29.1 | 30 | 39.5 | 4.2 | 39.4 | 29.0 | 49.5 |
| Self-reported health | Bogota | 184 | 64.7 | 19.4 | 68.8 | 12.5 | 100.0 | 170 | 72.6 | 17.4 | 75.0 | 0.0 | 100.0 |
| | Me'Phaa | 24 | 50.8 | 10.5 | 50.0 | 31.2 | 68.8 | 39 | 50.3 | 9.2 | 50.0 | 25.0 | 75.0 |
| | Mexico City | 30 | 56.0 | 7.6 | 56.2 | 43.8 | 75.0 | 30 | 60.4 | 8.6 | 62.5 | 37.5 | 75.0 |
| Visceral fat | Bogota | 184 | 3.9 | 1.3 | 4.0 | 1.0 | 8.0 | 170 | 5.4 | 2.8 | 5.0 | 1.0 | 14.0 |
| | Me'Phaa | 24 | 6.4 | 1.8 | 6.0 | 3.0 | 11.0 | 35 | 9.4 | 4.7 | 8.0 | 2.0 | 23.0 |
| | Mexico City | 30 | 6.4 | 2.0 | 7.0 | 2.0 | 10.0 | 30 | 6.2 | 3.2 | 6.0 | 2.0 | 17.0 |
| Waist circumference (cm) | Bogota | 184 | 71.8 | 8.4 | 70.3 | 55.3 | 103.9 | 170 | 78.2 | 7.9 | 77.6 | 62.1 | 103.6 |
| | Me'Phaa | 24 | 87.0 | 8.2 | 86.7 | 73.0 | 106.0 | 39 | 88.6 | 11.9 | 86.4 | 70.5 | 118.0 |
| | Mexico City | 30 | 87.8 | 10.9 | 87.4 | 66.5 | 113.9 | 30 | 84.5 | 8.4 | 84.3 | 69.0 | 106.6 |
| Weight (kg) | Bogota | 184 | 57.8 | 10.2 | 55.8 | 39.3 | 93.9 | 170 | 68.2 | 10.5 | 67.4 | 46.5 | 106.6 |
| | Me'Phaa | 24 | 54.2 | 7.7 | 54.4 | 43.7 | 67.2 | 39 | 65.9 | 14.5 | 61.9 | 43.4 | 101.7 |
| | Mexico City | 30 | 65.5 | 12.5 | 65.0 | 41.8 | 100.3 | 30 | 71.0 | 11.5 | 69.0 | 48.7 | 114.1 |

##### 2.1.2 Figure 1. Distribution by Sex, Population and Country

Kernel density plot for all measured variables by `Sample` (`Population` and `Country` interaction) and `Sex`.

```
datp <- melt(dat,
             id.vars = c(1:4, 15),
             measure.vars = 5:14,
             variable.name = "Measure",
             value.name = "Value")

levels(datp$Measure) <- c("Age",
                          "'Waist circumference (cm)'",
                          "'Hip (cm)'",
                          "'Height (cm)'",
                          "'Weight (kg)'",
                          "'Fat (%)'",
                          "'Visceral fat'",
```

```
                          "BMI~(kg/m^2)",
                          "'Muscle (%)'",
                          "'Self-reported health'")

datpF <- subset(datp, datp$Sex == "Women")
datpM <- subset(datp, datp$Sex == "Men")

colfunc <- colorRampPalette(c("deepskyblue2", "brown2")) # custom colour palette

Fig1a <- ggplot(datpF, aes(Value,
                           fill = Sample,
                           colour = Sample)) +
  geom_density(alpha = 0.3) +
  scale_fill_manual(values = colfunc(3)) +
  scale_color_manual(values = colfunc(3)) +
  facet_wrap(~Measure,
             scales = "free",
             ncol = 5,
             labeller = label_parsed) +
  labs(y = "Density",
       x = NULL,
       subtitle = "Women") +
  theme_light() +
  theme(strip.text.x = element_text(colour = "black"))

Fig1b <- ggplot(datpM, aes(Value,
                           fill = Sample,
                           colour = Sample)) +
  geom_density(alpha = 0.3) +
  scale_fill_manual(values = colfunc(3)) +
  scale_color_manual(values = colfunc(3)) +
  facet_wrap(. ~ Measure,
             scales = "free",
             ncol = 5,
             labeller = label_parsed) +
  labs(y = "Density",
       x = NULL,
       subtitle = "Men") +
  theme_light() +
  theme(strip.text.x = element_text(colour = "black"))

Fig1 <- ggarrange(Fig1a,
                  Fig1b,
                  nrow = 2,
                  ncol = 1,
                  labels = "auto",
                  legend = "bottom",
                  common.legend = TRUE)
Fig1
```

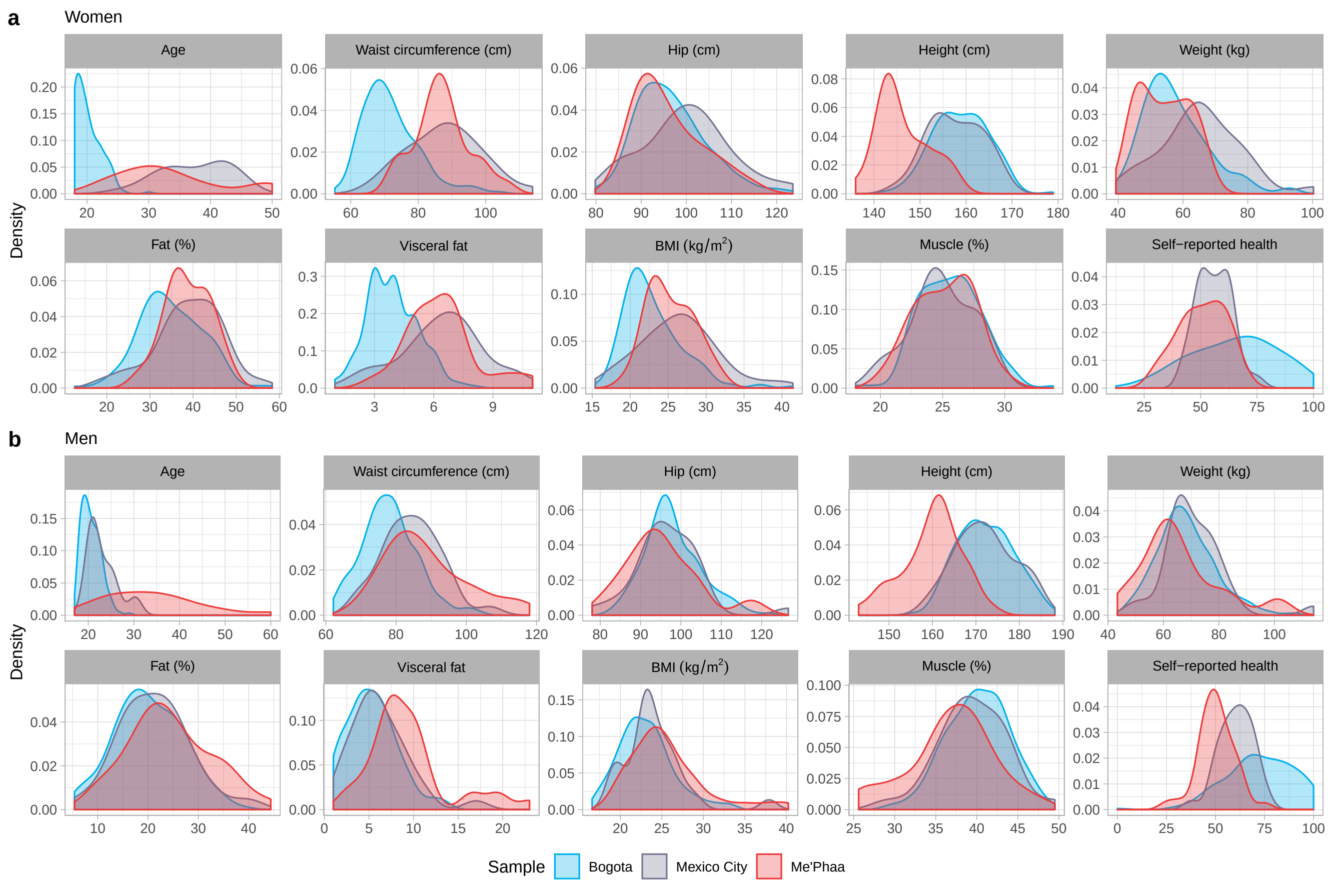

**Figure 1. Distribution of all measured variables by sex and sample (a)** Women. **(b)** Men.

##### 2.1.3 Figure S1. Health, height, and waist by Sex and Sample.

Comparison of height, waist and health between women and men from each sample.

```
t_testHea <- dat %>%
  group_by(Sample) %>%
  t_test(Health ~ Sex) %>%
  adjust_pvalue() %>%
  add_significance("p.adj")
t_testHea$p.adj.signif[t_testHea$p.adj.signif == "ns"] <- NA

FigS1a <- ggboxplot(dat,
  x = "Sex",
  y = "Health",
  color = "Sex",
  palette = rev(colfunc(2)),
  add = "jitter") +
  facet_wrap(~Sample) +
  stat_pvalue_manual(t_testHea,
                     label = "p.adj.signif",
                     y.position = 105,
                     tip.length = 0.01) +
  labs(x = NULL,
       y = "Self-reported health") +
  theme_light() +
  theme(strip.text.x = element_text(colour = "black"),
```

```
        legend.position = "none")

t_testHei <- dat %>%
  group_by(Sample) %>%
  t_test(Height ~ Sex) %>%
  adjust_pvalue() %>%
  add_significance("p.adj")
t_testHei$p.adj.signif[t_testHei$p.adj.signif == "ns"] <- NA

FigS1b <- ggboxplot(dat,
  x = "Sex",
  y = "Height",
  color = "Sex",
  palette = rev(colfunc(2)),
  add = "jitter") +
  facet_wrap(~Sample) +
  theme(legend.position = "none") +
  stat_pvalue_manual(t_testHei,
                     label = "p.adj.signif",
                     y.position = 195,
                     tip.length = 0.01) +
  labs(x = NULL,
       y = "Height (cm)") +
  theme_light() +
  theme(strip.text.x = element_text(colour = "black"))

t_testWai <- dat %>%
  group_by(Sample) %>%
  t_test(Waist ~ Sex) %>%
  adjust_pvalue() %>%
  add_significance("p.adj")
t_testWai$p.adj.signif[t_testWai$p.adj.signif == "ns"] <- NA

FigS1c <- ggboxplot(dat,
  x = "Sex",
  y = "Waist",
  color = "Sex",
  palette = rev(colfunc(2)),
  add = "jitter") +
  facet_wrap(~Sample) +
  theme(legend.position = "none") +
  stat_pvalue_manual(t_testWai,
                     label = "p.adj.signif",
                     y.position = 115,
                     tip.length = 0.01) +
  labs(x = NULL,
       y = "Waist circumference (cm)") +
  theme_light() +
  theme(strip.text.x = element_text(colour = "black"))

FigS1 <- ggarrange(FigS1a,
                   ggarrange(FigS1b,
                             FigS1c,
                             ncol = 2,
                             nrow = 1,
```

```
                      common.legend = TRUE,
                      legend = "bottom",
                      labels = c("b", "c")),
              nrow = 2,
              labels = "a",
              ncol = 1)
FigS1
```

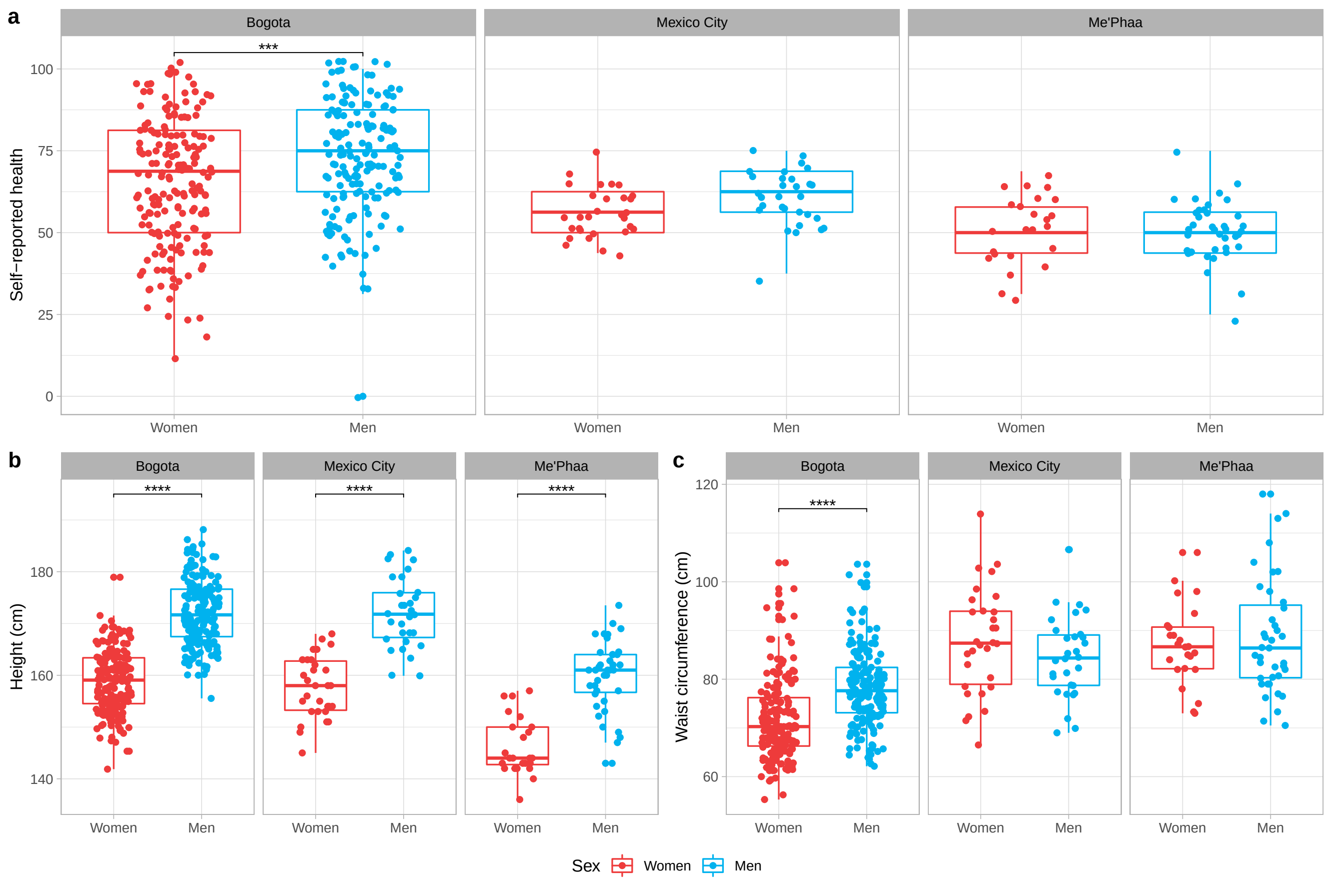

**Figure S1. Sexual dimorphism of height, waist and health for all samples** (**a**) Self-perceived health. (**b**) Height. (**c**) Waist. Comparisons between women and men for each sample, were performed using *t*-tests, adjusted for multiple tests. *** $p < 0.001$, **** $p < 0.0001$.

#### 2.2 Correlations

##### 2.2.1 Table S2. Correlation matrix (all participants)

```
corAll <- corstarsl(dat[, 5:14])
rownames(corAll) <- varnames
colnames(corAll) <- varnames[1:9]
TabS2 <- kable(
  corAll,
  booktabs = TRUE,
  align = "c",
  digits = 2,
  caption = "\\textbf{Table S2.} Correlations between measured variables
  for all participants",
  escape = FALSE) %>%
```

```
  kable_styling(latex_options = c("scale_down", "HOLD_position")) %>%
  footnote(
    general = "*$p$ < 0.05, **$p$ < 0.01, ***$p$ < 0.001",
    threeparttable = TRUE,
    escape = FALSE)
TabS2
```

**Table S2.** Correlations between measured variables for all participants

| | Age | Waist circumference (cm) | Hip (cm) | Height (cm) | Weight (kg) | Fat (%) | Visceral fat | BMI (kg/m$^2$) | Muscle (%) |
|---|---|---|---|---|---|---|---|---|---|
| Age | | | | | | | | | |
| Waist circumference (cm) | 0.46*** | | | | | | | | |
| Hip (cm) | 0.05 | 0.67*** | | | | | | | |
| Height (cm) | -0.29*** | 0.05 | 0.19*** | | | | | | |
| Weight (kg) | 0.03 | 0.73*** | 0.81*** | 0.50*** | | | | | |
| Fat (%) | 0.19*** | 0.37*** | 0.54*** | -0.55*** | 0.21*** | | | | |
| Visceral fat | 0.40*** | 0.79*** | 0.59*** | -0.06 | 0.68*** | 0.33*** | | | |
| BMI (kg/m$^2$) | 0.26*** | 0.81*** | 0.80*** | -0.14** | 0.78*** | 0.64*** | 0.80*** | | |
| Muscle (%) | -0.15*** | -0.06 | -0.25*** | 0.64*** | 0.13** | -0.91*** | -0.07 | -0.31*** | |
| Self-reported health | -0.24*** | -0.23*** | -0.10* | 0.27*** | -0.03 | -0.25*** | -0.20*** | -0.21*** | 0.23*** |

*Note:*
*$p < 0.05$, **$p < 0.01$, ***$p < 0.001$

##### 2.2.2 Table S3. Correlation matrix (women)

```
corF <- corstarsl(datF[, 5:14])
rownames(corF) <- varnames
colnames(corF) <- varnames[1:9]
TabS3 <- kable(
  corF,
  booktabs = TRUE, align = "c", digits = 2,
  caption = "\\textbf{Table S3.} Correlations between measured variables
  for women",
  escape = FALSE) %>%
  kable_styling(latex_options = c("scale_down", "HOLD_position")) %>%
  footnote(
    general = "*$p$ < 0.05, **$p$ < 0.01, ***$p$ < 0.001",
    threeparttable = TRUE,
    escape = FALSE)
TabS3
```

**Table S3.** Correlations between measured variables for women

| | Age | Waist circumference (cm) | Hip (cm) | Height (cm) | Weight (kg) | Fat (%) | Visceral fat | BMI (kg/m$^2$) | Muscle (%) |
|---|---|---|---|---|---|---|---|---|---|
| Age | | | | | | | | | |
| Waist circumference (cm) | 0.56*** | | | | | | | | |
| Hip (cm) | 0.13* | 0.69*** | | | | | | | |
| Height (cm) | -0.27*** | -0.19** | 0.15* | | | | | | |
| Weight (kg) | 0.12 | 0.73*** | 0.88*** | 0.29*** | | | | | |
| Fat (%) | 0.21** | 0.79*** | 0.84*** | -0.21** | 0.81*** | | | | |
| Visceral fat | 0.61*** | 0.87*** | 0.69*** | -0.36*** | 0.66*** | 0.84*** | | | |
| BMI (kg/m$^2$) | 0.26*** | 0.84*** | 0.83*** | -0.21** | 0.87*** | 0.93*** | 0.86*** | | |
| Muscle (%) | -0.12 | -0.54*** | -0.60*** | 0.28*** | -0.50*** | -0.81*** | -0.60*** | -0.65*** | |
| Self-reported health | -0.20** | -0.24*** | -0.11 | 0.23*** | -0.09 | -0.22*** | -0.25*** | -0.20** | 0.19** |

*Note:*
*$p < 0.05$, **$p < 0.01$, ***$p < 0.001$

##### 2.2.3 Table S4. Correlation matrix (men)

```
corM <- corstarsl(datM[, 5:14])
rownames(corM) <- varnames
```

```
colnames(corM) <- varnames[1:9]
TabS4 <- kable(
  corM,
  booktabs = TRUE,
  align = "c",
  digits = 2,
  caption = "\\textbf{Table S4.} Correlations between measured variables
  for men",
  escape = FALSE) %>%
  kable_styling(latex_options = c("scale_down", "HOLD_position")) %>%
  footnote(
    general = "*$p$ < 0.05, **$p$ < 0.01, ***$p$ < 0.001",
    threeparttable = TRUE,
    escape = FALSE)
TabS4
```

**Table S4.** Correlations between measured variables for men

| | Age | Waist circumference (cm) | Hip (cm) | Height (cm) | Weight (kg) | Fat (%) | Visceral fat | BMI (kg/m$^2$) | Muscle (%) |
|---|---|---|---|---|---|---|---|---|---|
| Age | | | | | | | | | |
| Waist circumference (cm) | 0.40*** | | | | | | | | |
| Hip (cm) | -0.05 | 0.68*** | | | | | | | |
| Height (cm) | -0.44*** | -0.13* | 0.31*** | | | | | | |
| Weight (kg) | -0.01 | 0.70*** | 0.87*** | 0.39*** | | | | | |
| Fat (%) | 0.26*** | 0.83*** | 0.74*** | -0.12 | 0.77*** | | | | |
| Visceral fat | 0.39*** | 0.83*** | 0.62*** | -0.34*** | 0.68*** | 0.86*** | | | |
| BMI (kg/m$^2$) | 0.26*** | 0.84*** | 0.76*** | -0.16* | 0.84*** | 0.90*** | 0.94*** | | |
| Muscle (%) | -0.37*** | -0.82*** | -0.69*** | 0.09 | -0.68*** | -0.93*** | -0.82*** | -0.79*** | |
| Self-reported health | -0.29*** | -0.32*** | -0.09 | 0.23*** | -0.10 | -0.21** | -0.29*** | -0.23*** | 0.24*** |

*Note:*
*$p < 0.05$, **$p < 0.01$, ***$p < 0.001$

### 3 Models to predict self-reported health

Given that there are missing data (NAs) on some variables on a few participants, and to ensure models were comparable (by AICc), participants with missing data (NAs) were excluded to always have the same $n$ regardless of which predictor variables are included.

In addition, because in interactions `Waist` and `Height` were included, these variables were mean-centred.

```
data <- dat[complete.cases(dat), ]
data$Age <- as.numeric(data$Age)
data$Waist_c <- c(scale(data$Waist, scale = FALSE))
data$Height_c <- c(scale(data$Height, scale = FALSE))
npersample <- data %>%
  group_by(Sample, Sex) %>%
  summarise(no_rows = length(Sex))
```

All models had a total $n$ of 473 (Bogota: 184 women and 170 men; Mexico City: 30 women and 30 men; Me'Phaa: 24 women and 35 men).

#### 3.1 Model fitting

We created three models.

##### 3.1.1 Model 1

The first, full model (Model 1; `mod1`), included `Age`, `Hip`, `Fat`, `VisceralFat`, `Weight`, `Muscle`, and `BMI` as main effects, as well as all main effects and possible interactions between any combination of `Height` (centred), `Waist` (centred), `Sex` and `Sample`, as predictors of `Health`.

###### 3.1.1.1 Table S5. Model 1 summary

```
mod1 <- lm(Health ~
             Sex * Waist_c * Height_c * Sample + Age +
             Hip + Fat + VisceralFat + Weight + Muscle + BMI,
           data = data)

summCols <- c(
  "$B$",
  "$SE(B)$",
  "95\\% CI",
  "$t$",
  "$p$")

ci.mod1 <- as.data.frame(confint(mod1))
ci.mod1$CI <- paste(round(ci.mod1$`2.5 %`, 3), round(ci.mod1$`97.5 %`, 3), sep = " - ")
s.mod1 <- summary(mod1)
tabm1 <- as.data.frame(s.mod1$coefficients)
tabm1 <- cbind(tabm1, ci.mod1$CI)
tabm1 <- summaSig(tabm1, 4)
tabm1 <- summTerms(tabm1)
tabm1 <- tabm1[,c(1,2,5,3,4)]
colnames(tabm1) <- summCols

TabS5 <- kable(
  tabm1,
  digits = 2,
  booktabs = TRUE,
  align = "c",
  caption = "\\textbf{Table S5.} Model 1 Summary",
  escape = FALSE) %>%
  kable_styling(latex_options = "HOLD_position") %>%
  footnote(general = paste0(
    "$R^2$ = ",
    round(s.mod1$r.squared, 3),
    ", $R^2_{adjusted}$ = ",
    round(s.mod1$adj.r.squared, 3),
    ", $F$(",
    paste(s.mod1$fstatistic[2],
          s.mod1$fstatistic[3], sep = ", "),
    ") = ", round(s.mod1$fstatistic[1], 2),
    ", $p$ ",
    pvalr(pf(s.mod1$fstatistic[1],
             s.mod1$fstatistic[2],
             s.mod1$fstatistic[3],
             lower.tail = FALSE),
          digits = 4),
    ". Women and Bogota were used as reference categories for Sex and
    Sample, respectively. For model terms: WC(c) = Waist circumference (centred);
    H(c) = Height (centred); S = Sample. Significant predictors are in bold."),
    escape = FALSE,
    threeparttable = TRUE)
TabS5
```

**Table S5.** Model 1 Summary

| | $B$ | $SE(B)$ | 95% CI | $t$ | $p$ |
|---|---|---|---|---|---|
| (Intercept) | 78.72 | 31.30 | 17.206 — 140.23 | 2.52 | **0.012** |
| Sex(men) | -0.22 | 8.24 | -16.412 — 15.97 | -0.03 | 0.979 |
| WC(c) | -0.17 | 0.29 | -0.733 — 0.398 | -0.58 | 0.561 |
| H(c) | 0.67 | 0.64 | -0.597 — 1.931 | 1.04 | 0.3 |
| S(Mexico City) | -10.28 | 7.10 | -24.24 — 3.675 | -1.45 | 0.148 |
| S(Me'Phaa) | -46.92 | 23.66 | -93.422 — -0.419 | -1.98 | **0.048** |
| Age | 0.19 | 0.22 | -0.243 — 0.618 | 0.86 | 0.393 |
| Hip | -0.31 | 0.24 | -0.774 — 0.157 | -1.30 | 0.193 |
| Fat | -0.06 | 0.47 | -0.984 — 0.859 | -0.13 | 0.894 |
| Visceral Fat | 0.05 | 1.00 | -1.914 — 2.021 | 0.05 | 0.957 |
| Weight | -0.35 | 0.71 | -1.747 — 1.05 | -0.49 | 0.625 |
| Muscle | 0.38 | 0.67 | -0.94 — 1.7 | 0.57 | 0.572 |
| BMI | 1.18 | 1.88 | -2.518 — 4.888 | 0.63 | 0.53 |
| Sex(men) × WC(c) | 0.51 | 0.40 | -0.262 — 1.29 | 1.30 | 0.194 |
| Sex(men) × H(c) | -0.34 | 0.34 | -1.011 — 0.322 | -1.02 | 0.31 |
| WC(c) × H(c) | -0.01 | 0.03 | -0.064 — 0.034 | -0.60 | 0.55 |
| Sex(men) × S(Mexico City) | -4.57 | 9.27 | -22.784 — 13.649 | -0.49 | 0.622 |
| Sex(men) × S(Me'Phaa) | 20.01 | 24.14 | -27.435 — 67.449 | 0.83 | 0.408 |
| WC(c) × S(Mexico City) | 0.26 | 0.48 | -0.678 — 1.205 | 0.55 | 0.582 |
| WC(c) × S(Me'Phaa) | 2.55 | 2.03 | -1.426 — 6.535 | 1.26 | 0.208 |
| H(c) × S(Mexico City) | 0.17 | 0.73 | -1.262 — 1.6 | 0.23 | 0.817 |
| H(c) × S(Me'Phaa) | -1.92 | 1.20 | -4.266 — 0.434 | -1.60 | 0.11 |
| Sex(men) × WC(c) × H(c) | -0.03 | 0.03 | -0.096 — 0.04 | -0.82 | 0.413 |
| Sex(men) × WC(c) × S(Mexico City) | -0.89 | 0.79 | -2.44 — 0.653 | -1.14 | 0.257 |
| Sex(men) × WC(c) × S(Me'Phaa) | -2.71 | 2.07 | -6.782 — 1.36 | -1.31 | 0.191 |
| Sex(men) × H(c) × S(Mexico City) | 0.04 | 0.92 | -1.77 — 1.852 | 0.04 | 0.964 |
| Sex(men) × H(c) × S(Me'Phaa) | 1.60 | 1.40 | -1.144 — 4.348 | 1.15 | 0.252 |
| WC(c) × H(c) × S(Mexico City) | 0.02 | 0.06 | -0.103 — 0.138 | 0.28 | 0.778 |
| WC(c) × H(c) × S(Me'Phaa) | 0.15 | 0.11 | -0.068 — 0.366 | 1.35 | 0.178 |
| Sex(men) × WC(c) × H(c) × S(Mexico City) | 0.07 | 0.09 | -0.117 — 0.255 | 0.73 | 0.464 |
| Sex(men) × WC(c) × H(c) × S(Me'Phaa) | -0.11 | 0.12 | -0.355 — 0.132 | -0.90 | 0.368 |

*Note:*
$R^2 = 0.21$, $R^2_{adjusted} = 0.156$, $F(30, 442) = 3.91$, $p < 0.001$. Women and Bogota were used as reference categories for Sex and Sample, respectively. For model terms: WC(c) = Waist circumference (centred); H(c) = Height (centred); S = Sample. Significant predictors are in bold.

##### 3.1.2 Model 2

To increase parsimony, two additional models were fitted. Model 2 (`mod2`), included only `Age`, and main effects and all possible interactions between any combination of `Waist` (centred), `Height` (centred), `Sex`, and `Sample` as predictors of `Health`.

###### 3.1.2.1 Table S6. Model 2 summary

```
mod2 <- lm(Health ~
Sex * Waist_c * Height_c * Sample + Age,
data = data,
na.action = "na.fail")

ci.mod2 <- as.data.frame(confint(mod2))
```

```
ci.mod2$CI <- paste(round(ci.mod2$`2.5 %`, 3), round(ci.mod2$`97.5 %`, 3), sep = " - ")
s.mod2 <- summary(mod2)
tabm2 <- as.data.frame(s.mod2$coefficients)
tabm2 <- cbind(tabm2, ci.mod2$CI)
tabm2 <- summaSig(tabm2, 4)
tabm2 <- summTerms(tabm2)
tabm2 <- tabm2[,c(1,2,5,3,4)]
colnames(tabm2) <- summCols

TabS6 <- kable(
  tabm2,
  digits = 2,
  booktabs = TRUE,
  align = "c",
  caption = "\\textbf{Table S6.} Model 2 Summary",
  escape = FALSE) %>%
  kable_styling(latex_options = "HOLD_position") %>%
  footnote(general = paste0(
    "$R^2$ = ",
    round(s.mod2$r.squared, 3),
    ", $R^2_{adjusted}$ = ",
    round(s.mod2$adj.r.squared, 3),
    ", $F$(",
    paste(s.mod2$fstatistic[2],
          s.mod2$fstatistic[3], sep = ", "),
    ") = ", round(s.mod2$fstatistic[1], 2),
    ", $p$ ",
    pvalr(pf(s.mod2$fstatistic[1],
             s.mod2$fstatistic[2],
             s.mod2$fstatistic[3],
             lower.tail = FALSE),
          digits = 4),
    ". Women and Bogota were used as reference categories for Sex and
    Sample, respectively. For model terms: WC(c) = Waist circumference (centred);
    H(c) = Height (centred); S = Sample. Significant predictors are in bold."),
    escape = FALSE,
    threeparttable = TRUE)
TabS6
```

**Table S6.** Model 2 Summary

| | $B$ | $SE(B)$ | 95% CI | $t$ | $p$ |
|---|---|---|---|---|---|
| (Intercept) | 61.13 | 4.57 | 52.152 — 70.105 | 13.38 | **<0.0001** |
| Sex(men) | 8.43 | 2.83 | 2.865 — 14 | 2.98 | **0.003** |
| WC(c) | -0.42 | 0.20 | -0.805 — -0.031 | -2.12 | **0.034** |
| H(c) | 0.36 | 0.24 | -0.104 — 0.821 | 1.52 | 0.129 |
| S(Mexico City) | -7.75 | 6.91 | -21.322 — 5.822 | -1.12 | 0.262 |
| S(Me'Phaa) | -41.88 | 23.40 | -87.868 — 4.113 | -1.79 | 0.074 |
| Age | 0.17 | 0.20 | -0.227 — 0.576 | 0.85 | 0.393 |
| Sex(men) × WC(c) | 0.41 | 0.33 | -0.238 — 1.063 | 1.25 | 0.213 |
| Sex(men) × H(c) | -0.40 | 0.31 | -1.007 — 0.202 | -1.31 | 0.191 |
| WC(c) × H(c) | -0.02 | 0.02 | -0.065 — 0.023 | -0.94 | 0.349 |
| Sex(men) × S(Mexico City) | -4.72 | 9.17 | -22.749 — 13.31 | -0.51 | 0.607 |
| Sex(men) × S(Me'Phaa) | 17.40 | 23.96 | -29.689 — 64.487 | 0.73 | 0.468 |
| WC(c) × S(Mexico City) | 0.31 | 0.48 | -0.629 — 1.242 | 0.64 | 0.521 |
| WC(c) × S(Me'Phaa) | 2.51 | 2.02 | -1.455 — 6.482 | 1.24 | 0.214 |
| H(c) × S(Mexico City) | 0.14 | 0.72 | -1.281 — 1.556 | 0.19 | 0.849 |
| H(c) × S(Me'Phaa) | -1.80 | 1.19 | -4.131 — 0.534 | -1.52 | 0.13 |
| Sex(men) × WC(c) × H(c) | -0.02 | 0.03 | -0.089 — 0.043 | -0.68 | 0.496 |
| Sex(men) × WC(c) × S(Mexico City) | -0.78 | 0.78 | -2.304 — 0.753 | -1.00 | 0.319 |
| Sex(men) × WC(c) × S(Me'Phaa) | -2.58 | 2.06 | -6.641 — 1.473 | -1.25 | 0.211 |
| Sex(men) × H(c) × S(Mexico City) | 0.03 | 0.92 | -1.769 — 1.837 | 0.04 | 0.97 |
| Sex(men) × H(c) × S(Me'Phaa) | 1.64 | 1.39 | -1.087 — 4.372 | 1.18 | 0.238 |
| WC(c) × H(c) × S(Mexico City) | 0.02 | 0.06 | -0.104 — 0.137 | 0.27 | 0.788 |
| WC(c) × H(c) × S(Me'Phaa) | 0.15 | 0.11 | -0.069 — 0.363 | 1.33 | 0.183 |
| Sex(men) × WC(c) × H(c) × S(Mexico City) | 0.06 | 0.09 | -0.129 — 0.24 | 0.59 | 0.553 |
| Sex(men) × WC(c) × H(c) × S(Me'Phaa) | -0.11 | 0.12 | -0.353 — 0.131 | -0.90 | 0.368 |

*Note:*
$R^2 = 0.201$, $R^2_{adjusted} = 0.158$, $F(24, 448) = 4.7$, $p < 0.001$. Women and Bogota were used as reference categories for Sex and Sample, respectively. For model terms: WC(c) = Waist circumference (centred); H(c) = Height (centred); S = Sample. Significant predictors are in bold.

##### 3.1.3 Model 3

Finally, for Model 3 (`mod3`), we used the functions `dredge` and `model.sel` from the package `MuMIn` (Multi-Model Inference); the first function creates a set of models with combinations (subsets) of fixed effect terms, from Model 2 (`mod2`), and the second builds a model selection table. In our case, these functions created and compared 334 models.

```
fitt <- dredge(mod2)
options(digits = 2)
m.sel <- model.sel(fitt)
```

###### 3.1.3.1 Figure 2. Model selection plot

The best model (labelled 159, Fig. S2), included `Height` (centred), `Sample`, `Sex`, `Waist` (centred) and the interaction between `Height` (centred) and `Waist` (centred). However, given the age differences between samples, we selected the second-best model (labelled 160, Fig. S2), because it also included `Age` as a regressor, and had a $\Delta AICc$ of less than 2 compared to the best model. This model, including `Age`, was therefore selected as our final model (Model 3).

```
par(mar=c(1,4,10,3))
plot(fitt, labels = c(
  "Intercept",
```

```
  "Age",
  "H(c)",
  "S",
  "Sex",
  "WC(c)",
  "H(c):S",
  "H(c):Sex",
  "H(c):WC(c)",
  "S:Sex",
  "S:WC(c)",
  "Sex:WC(c)",
  "H(c):S:Sex",
  "H(c):S:WC(c)",
  "H(c):Sex:WC(c)",
  "S:Sex:WC(c)",
  "H(c):S:Sex:WC(c)"),
  ylab = expression("Cumulative " ~italic(w[i]*(AICc))),
  col = c(colfunc(1)), border = "gray30",
  labAsExpr = TRUE)
```

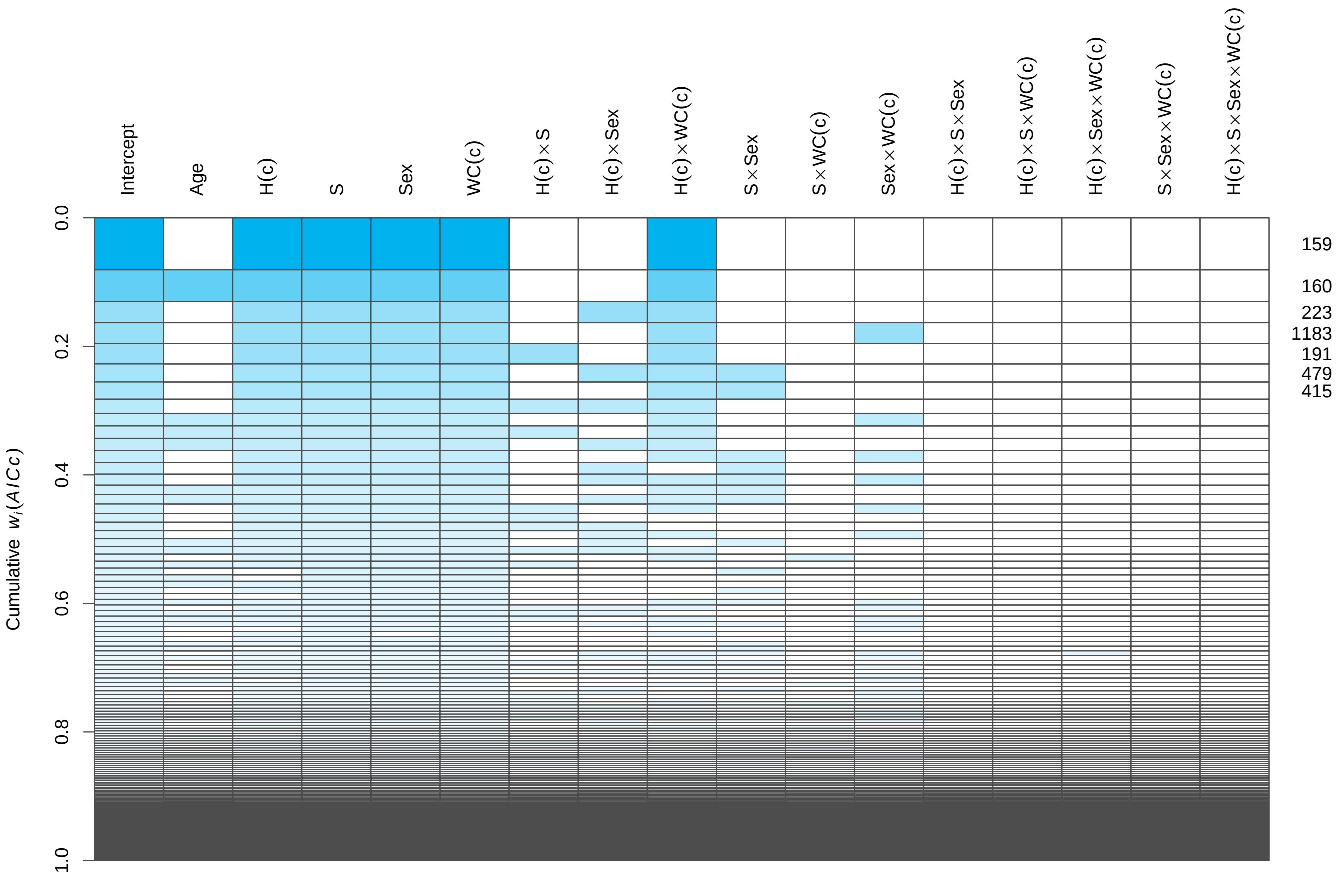

**Figure 2. Model selection plot.** Rows represent each of the 334 compared models. Cells coloured in blue represent included terms in each model, according to their Akaike weight ($w_i(AICc)$), represented as the height of each row/model. Given the important age differences between samples, we selected the second-best model (labelled **160**), because it had the same structure as the best model (labelled **159**), but also included Age as a regressor. Furthermore, this second-best model had a $\Delta AICc$ of less than 2 units ($\approx 0.98$) compared to the best model. For model terms: WC(c) = Waist circumference (centred); H(c) = Height (centred); S = Sample.

###### 3.1.3.2 Table S7. Model 3 summary

```
# Select second-best model and assign it as Model 3
mod3 <- get.models(fitt, subset = 2)[[1]]
s.mod3 <- summary(mod3)
ci.mod3 <- as.data.frame(confint(mod3))
ci.mod3$CI <- paste(round(ci.mod3$`2.5 %`, 3), round(ci.mod3$`97.5 %`, 3), sep = " - ")
tabm3 <- as.data.frame(s.mod3$coefficients)
tabm3 <- cbind(tabm3, ci.mod3$CI)
tabm3 <- summaSig(tabm3, 4)
tabm3 <- summTerms(tabm3)
tabm3 <- tabm3[,c(1,2,5,3,4)]
colnames(tabm3) <- summCols

TabS7 <- kable(tabm3, digits = 2,
  booktabs = TRUE, align = "c",
  caption = "\\textbf{Table S7.} Model 3 Summary",
  escape = FALSE) %>%
  kable_styling(latex_options = "HOLD_position") %>%
  footnote(general = paste0(
    "$R^2$ = ", round(s.mod3$r.squared, 3),
    ", $R^2_{adjusted}$ = ",
    round(s.mod3$adj.r.squared, 3), ", $F$(",
    paste(s.mod3$fstatistic[2],
          s.mod3$fstatistic[3], sep = ", "),
    ") = ", round(s.mod3$fstatistic[1], 2),
    ", $p$ ",
    pvalr(pf(s.mod3$fstatistic[1],
             s.mod3$fstatistic[2],
             s.mod3$fstatistic[3],
             lower.tail = FALSE),
          digits = 4),
    ". Women and Bogota were used as reference categories for Sex and
    Sample, respectively. For model terms: WC(c) = Waist circumference (centred);
    H(c) = Height (centred); S = Sample. Significant predictors are in bold."),
    escape = FALSE, threeparttable = TRUE)
TabS7
```

**Table S7.** Model 3 Summary

| | $B$ | $SE(B)$ | 95% CI | $t$ | $p$ |
|---|---|---|---|---|---|
| (Intercept) | 61.40 | 3.77 | 53.983 — 68.815 | 16.3 | **<0.0001** |
| Age | 0.17 | 0.16 | -0.153 — 0.495 | 1.0 | 0.299 |
| H(c) | 0.16 | 0.12 | -0.077 — 0.403 | 1.3 | 0.183 |
| S(Mexico City) | -9.32 | 2.87 | -14.96 — -3.681 | -3.2 | **0.001** |
| S(Me'Phaa) | -16.78 | 3.53 | -23.722 — -9.83 | -4.8 | **<0.0001** |
| Sex(men) | 6.01 | 2.26 | 1.564 — 10.458 | 2.7 | **0.008** |
| WC(c) | -0.28 | 0.09 | -0.46 — -0.11 | -3.2 | **0.001** |
| H(c) × WC(c) | -0.02 | 0.01 | -0.036 — -0.003 | -2.3 | **0.022** |

*Note:*
$R^2$ = 0.183, $R^2_{adjusted}$ = 0.171, $F$(7, 465) = 14.88, $p$ < 0.001. Women and Bogota were used as reference categories for Sex and Sample, respectively. For model terms: WC(c) = Waist circumference (centred); H(c) = Height (centred); S = Sample. Significant predictors are in bold.

#### 3.2 Model comparison

##### 3.2.1 Table 2. Summary of the three models

```
tab1_2 <- merge(tabm1[,c(1,2,4,5)],tabm2[,c(1,2,4,5)], by= "row.names", all = TRUE)
rownames(tab1_2) <- tab1_2[,1]
tab1_2[,1] <- NULL
#reorder the interaction terms to match the other model summaries (first WC and then H)
rownames(tabm3)[rownames(tabm3) ==
                  "H(c) $\\times$ WC(c)"] <-
  "WC(c) $\\times$ H(c)"
tab1_3 <- merge(tab1_2,tabm3[,c(1,2,4,5)], by= "row.names", all = TRUE)
rownames(tab1_3) <- tab1_3[,1]
tab1_3[,1] <- NULL

Tab2 <- kable(
  tab1_3,
  digits = 2,
  booktabs = TRUE,
  align = "c",
  caption = "\\textbf{Table 2.} Results of separate LMs testing effects of independent
  variables on self-reported health",
  col.names = rep(summCols[c(1,2,4,5)], 3),
  escape = FALSE) %>%
  add_header_above(c(" " = 1,
                     "Model 1" = 4,
                     "Model 2" = 4,
                     "Model 3" = 4)) %>%
  kable_styling(latex_options = c("scale_down", "HOLD_position")) %>%
  footnote(general = paste0(
    " For Model 1, $R^2$ = ",
    round(s.mod1$r.squared, 3),
  ", $R^2_{adjusted}$ = ",
  round(s.mod1$adj.r.squared, 3),
  ", $F$(",
  paste(s.mod1$fstatistic[2],
        s.mod1$fstatistic[3], sep = ", "),
  ") = ", round(s.mod1$fstatistic[1], 2),
  ", $p$ ",
  pvalr(pf(s.mod1$fstatistic[1],
           s.mod1$fstatistic[2],
           s.mod1$fstatistic[3],
           lower.tail = FALSE),
        digits = 4),
  "; for Model 2, $R^2$ = ",
  round(s.mod2$r.squared, 3),
  ", $R^2_{adjusted}$ = ",
  round(s.mod2$adj.r.squared, 3),
  ", $F$(",
  paste(s.mod2$fstatistic[2],
        s.mod2$fstatistic[3], sep = ", "),
  ") = ", round(s.mod2$fstatistic[1], 2),
  ", $p$ ",
  pvalr(pf(s.mod2$fstatistic[1],
           s.mod2$fstatistic[2],
           s.mod2$fstatistic[3],
           lower.tail = FALSE),
```

```
      digits = 4),
  "; for Model 3, $R^2$ = ",
  round(s.mod3$r.squared, 3),
  ", $R^2_{adjusted}$ = ",
  round(s.mod3$adj.r.squared, 3),
  ", $F$(",
  paste(s.mod3$fstatistic[2],
        s.mod3$fstatistic[3], sep = ", "),
  ") = ", round(s.mod3$fstatistic[1], 2),
  ", $p$ ",
  pvalr(pf(s.mod3$fstatistic[1],
           s.mod3$fstatistic[2],
           s.mod3$fstatistic[3],
           lower.tail = FALSE),
        digits = 4),
  ". Women and Bogota were used as reference categories
  for Sex and Sample, respectively. For model terms: WC(c) = Waist circumference
  (centred); H(c) = Height (centred); S = Sample. Significant predictors are in
  bold."),
  escape = FALSE,
  threeparttable = TRUE)
Tab2
```

**Table 2.** Results of separate LMs testing effects of independent variables on self-reported health

| | Model 1 | | | | Model 2 | | | | Model 3 | | | |
|---|---|---|---|---|---|---|---|---|---|---|---|---|
| | $B$ | $SE(B)$ | $t$ | $p$ | $B$ | $SE(B)$ | $t$ | $p$ | $B$ | $SE(B)$ | $t$ | $p$ |
| (Intercept) | 78.72 | 31.30 | 2.52 | **0.012** | 61.13 | 4.57 | 13.38 | **<0.0001** | 61.40 | 3.77 | 16.3 | **<0.0001** |
| Age | 0.19 | 0.22 | 0.86 | 0.393 | 0.17 | 0.20 | 0.85 | 0.393 | 0.17 | 0.16 | 1.0 | 0.299 |
| BMI | 1.18 | 1.88 | 0.63 | 0.53 | | | | | | | | |
| Fat | -0.06 | 0.47 | -0.13 | 0.894 | | | | | | | | |
| H(c) | 0.67 | 0.64 | 1.04 | 0.3 | 0.36 | 0.24 | 1.52 | 0.129 | 0.16 | 0.12 | 1.3 | 0.183 |
| H(c) × S(Me'Phaa) | -1.92 | 1.20 | -1.60 | 0.11 | -1.80 | 1.19 | -1.52 | 0.13 | | | | |
| H(c) × S(Mexico City) | 0.17 | 0.73 | 0.23 | 0.817 | 0.14 | 0.72 | 0.19 | 0.849 | | | | |
| Hip | -0.31 | 0.24 | -1.30 | 0.193 | | | | | | | | |
| Muscle | 0.38 | 0.67 | 0.57 | 0.572 | | | | | | | | |
| S(Me'Phaa) | -46.92 | 23.66 | -1.98 | **0.048** | -41.88 | 23.40 | -1.79 | 0.074 | -16.78 | 3.53 | -4.8 | **<0.0001** |
| S(Mexico City) | -10.28 | 7.10 | -1.45 | 0.148 | -7.75 | 6.91 | -1.12 | 0.262 | -9.32 | 2.87 | -3.2 | **0.001** |
| Sex(men) | -0.22 | 8.24 | -0.03 | 0.979 | 8.43 | 2.83 | 2.98 | **0.003** | 6.01 | 2.26 | 2.7 | **0.008** |
| Sex(men) × H(c) | -0.34 | 0.34 | -1.02 | 0.31 | -0.40 | 0.31 | -1.31 | 0.191 | | | | |
| Sex(men) × H(c) × S(Me'Phaa) | 1.60 | 1.40 | 1.15 | 0.252 | 1.64 | 1.39 | 1.18 | 0.238 | | | | |
| Sex(men) × H(c) × S(Mexico City) | 0.04 | 0.92 | 0.04 | 0.964 | 0.03 | 0.92 | 0.04 | 0.97 | | | | |
| Sex(men) × S(Me'Phaa) | 20.01 | 24.14 | 0.83 | 0.408 | 17.40 | 23.96 | 0.73 | 0.468 | | | | |
| Sex(men) × S(Mexico City) | -4.57 | 9.27 | -0.49 | 0.622 | -4.72 | 9.17 | -0.51 | 0.607 | | | | |
| Sex(men) × WC(c) | 0.51 | 0.40 | 1.30 | 0.194 | 0.41 | 0.33 | 1.25 | 0.213 | | | | |
| Sex(men) × WC(c) × H(c) | -0.03 | 0.03 | -0.82 | 0.413 | -0.02 | 0.03 | -0.68 | 0.496 | | | | |
| Sex(men) × WC(c) × H(c) × S(Me'Phaa) | -0.11 | 0.12 | -0.90 | 0.368 | -0.11 | 0.12 | -0.90 | 0.368 | | | | |
| Sex(men) × WC(c) × H(c) × S(Mexico City) | 0.07 | 0.09 | 0.73 | 0.464 | 0.06 | 0.09 | 0.59 | 0.553 | | | | |
| Sex(men) × WC(c) × S(Me'Phaa) | -2.71 | 2.07 | -1.31 | 0.191 | -2.58 | 2.06 | -1.25 | 0.211 | | | | |
| Sex(men) × WC(c) × S(Mexico City) | -0.89 | 0.79 | -1.14 | 0.257 | -0.78 | 0.78 | -1.00 | 0.319 | | | | |
| Visceral Fat | 0.05 | 1.00 | 0.05 | 0.957 | | | | | | | | |
| WC(c) | -0.17 | 0.29 | -0.58 | 0.561 | -0.42 | 0.20 | -2.12 | **0.034** | -0.28 | 0.09 | -3.2 | **0.001** |
| WC(c) × H(c) | -0.01 | 0.03 | -0.60 | 0.55 | -0.02 | 0.02 | -0.94 | 0.349 | -0.02 | 0.01 | -2.3 | **0.022** |
| WC(c) × H(c) × S(Me'Phaa) | 0.15 | 0.11 | 1.35 | 0.178 | 0.15 | 0.11 | 1.33 | 0.183 | | | | |
| WC(c) × H(c) × S(Mexico City) | 0.02 | 0.06 | 0.28 | 0.778 | 0.02 | 0.06 | 0.27 | 0.788 | | | | |
| WC(c) × S(Me'Phaa) | 2.55 | 2.03 | 1.26 | 0.208 | 2.51 | 2.02 | 1.24 | 0.214 | | | | |
| WC(c) × S(Mexico City) | 0.26 | 0.48 | 0.55 | 0.582 | 0.31 | 0.48 | 0.64 | 0.521 | | | | |
| Weight | -0.35 | 0.71 | -0.49 | 0.625 | | | | | | | | |

*Note:*
For Model 1, $R^2 = 0.21$, $R^2_{adjusted} = 0.156$, $F(30, 442) = 3.91$, $p < 0.001$; for Model 2, $R^2 = 0.201$, $R^2_{adjusted} = 0.158$, $F(24, 448) = 4.7$, $p < 0.001$; for Model 3, $R^2 = 0.183$, $R^2_{adjusted} = 0.171$, $F(7, 465) = 14.88$, $p < 0.001$. Women and Bogota were used as reference categories for Sex and Sample, respectively. For model terms: WC(c) = Waist circumference (centred); H(c) = Height (centred); S = Sample. Significant predictors are in bold.

##### 3.2.2 Table 3. Information criteria for the three models

In addition to being more parsimonious, Model 3 (mod3) had a lower Akaike information criterion ($AICc$), higher Akaike weight ($w_i(AICc)$), and higher $R^2_{adjusted}$ than the other two models.

```
aic1_3 <- AICctab(mod1,
                  mod2,
                  mod3,
                  weights = TRUE,
                  base = TRUE)
class(aic1_3) <- "data.frame"
tab3 <- aic1_3
row.names(tab3) <- c("Model 3", "Model 2", "Model 1")
tab3$weight <- format(round(tab3$weight, 8),
                       nsmall = 8,
                       scientific = FALSE)

Tab3 <- kable(
  tab3,
  align = "c",
  digits = 20,
  caption = "\\textbf{Table 3.} Information criteria for the three models",
  col.names = c("$AICc$", "$\\Delta AICc$", "$df$", "$w_{i}(AICc)$"),
  booktabs = TRUE,
  escape = FALSE) %>%
  kable_styling(latex_options = "HOLD_position") %>%
  footnote(general = paste0("Model 31 is close to ",
                            format(round(aic1_3[1,4]/aic1_3[2,4], 10),
                                   big.mark = ","),
                            " times more likely to be the best model
                            compared to Model 2, and about ",
                            format(round(aic1_3[1,4]/aic1_3[3,4], 12),
                                   big.mark = ",", scientific = FALSE),
                            " times compared to Model 1 (Model 2, was around ",
                            format(round(aic1_3[2,4]/aic1_3[3,4], 2),
                                   big.mark = ","), " times more likely
                            compared to Model 1). For a detailed description of values,
                            see the \\\\href{https://www.shorturl.at/iGIKT}{ICtab}
                            function documentation."),
           escape = FALSE,
           threeparttable = TRUE)
Tab3
```

**Table 3.** Information criteria for the three models

| | $AICc$ | $\Delta AICc$ | $df$ | $w_i(AICc)$ |
|---|---|---|---|---|
| Model 3 | 3999 | 0 | 9 | 0.99999782 |
| Model 2 | 4025 | 26 | 26 | 0.00000215 |
| Model 1 | 4034 | 35 | 32 | 0.00000003 |

*Note:*
Model 31 is close to 464,686 times more likely to be the best model compared to Model 2, and about 35,141,683 times compared to Model 1 (Model 2, was around 76 times more likely compared to Model 1). For a detailed description of values, see the ICtab function documentation.

#### 3.3 Final model (Model 3)

##### 3.3.1 Model diagnostics

###### 3.3.1.1 Table S8. Variance Inflation Factors of Information criteria for Final Model (Model 3) predictors

```
mod3VIF <- as.data.frame(vif(mod3))
row.names(mod3VIF) <- c("Age",
                        "Height(c)",
                        "Sample",
                        "Sex",
                        "Waist circumference(c)",
                        "Height(c) $\\times$ Waist(c)")

TabS8 <- kable(mod3VIF,
      booktabs = TRUE,
      digits = 2,
      align = "c",
      caption = "\\textbf{Table S8.} Variance Inflation Factors of the Final Model
      (Model 3) predictors",
      col.names = c("$GVIF$", "$df$", "$GVIF^{1/(2 \\times df)}$"),
      escape = FALSE) %>%
        kable_styling(latex_options = "HOLD_position") %>%
  footnote(general = "For a detailed description of values, see
  \\\\href{https://www.rdocumentation.org/packages/car/versions/3.0-5/topics/vif}{vif}
  function documentation.",
           escape = FALSE,
           threeparttable = TRUE)
TabS8
```

**Table S8.** Variance Inflation Factors of the Final Model (Model 3) predictors

| | $GVIF$ | $df$ | $GVIF^{1/(2\times df)}$ |
|---|---|---|---|
| Age | 2.4 | 1 | 1.5 |
| Height(c) | 2.5 | 1 | 1.6 |
| Sample | 3.0 | 2 | 1.3 |
| Sex | 2.2 | 1 | 1.5 |
| Waist circumference(c) | 1.5 | 1 | 1.2 |
| Height(c) $\times$ Waist(c) | 1.2 | 1 | 1.1 |

*Note:*
For a detailed description of values, see vif function documentation.

###### 3.3.1.2 Figure S2. Residual distribution by sample and linearity in each (single term) factor

```
FigS2a <- ggplot(data = mod3$model,
                 mapping = aes(sample = residuals(mod3))) +
  stat_qq_band(alpha = 0.3) +
  stat_qq_line(color = colfunc(2)[1]) +
  stat_qq_point(alpha = 0.3) +
  facet_wrap(~Sample, scales = "free") +
  labs(x = "Theoretical Quantiles",
       y = "Sample Quantiles") +
  theme_light() +
  theme(strip.text.x = element_text(colour = "black"))
```

```
FigS2b1 <- ggplot(data.frame(x1 = mod3$model$Waist_c,
                             pearson = residuals(mod3,
                                                 type = "pearson")),
                 aes(x = x1,
                     y = pearson)) +
  geom_point(alpha = 0.3) +
  geom_smooth(method = "lm",
              color = colfunc(2)[2]) +
  labs(x = "Waist circumference (centred)",
       y = "Pearson residuals") +
  theme_light()

FigS2b2 <- ggplot(data.frame(x1 = mod3$model$Height_c,
                             pearson = residuals(mod3,
                                                 type = "pearson")),
                 aes(x = x1,
                     y = pearson)) +
  geom_point(alpha = 0.3) +
  geom_smooth(method = "lm",
              color = colfunc(2)[2]) +
  labs(x = "Height (centred)",
       y = NULL) +
  theme_light()

FigS2b3 <- ggplot(data.frame(x1 = mod3$model$Age,
                             pearson = residuals(mod3,
                                                 type = "pearson")),
                 aes(x = x1,
                     y = pearson)) +
  geom_point(alpha = 0.3) +
  geom_smooth(method = "lm",
              color = colfunc(2)[2]) +
  labs(x = "Age",
       y = NULL) +
  theme_light()

FigS2 <- ggarrange(FigS2a,
                   ggarrange(FigS2b1,
                             FigS2b2,
                             FigS2b3,
                             ncol = 3),
                   nrow = 2,
                   labels = "auto")
FigS2
```

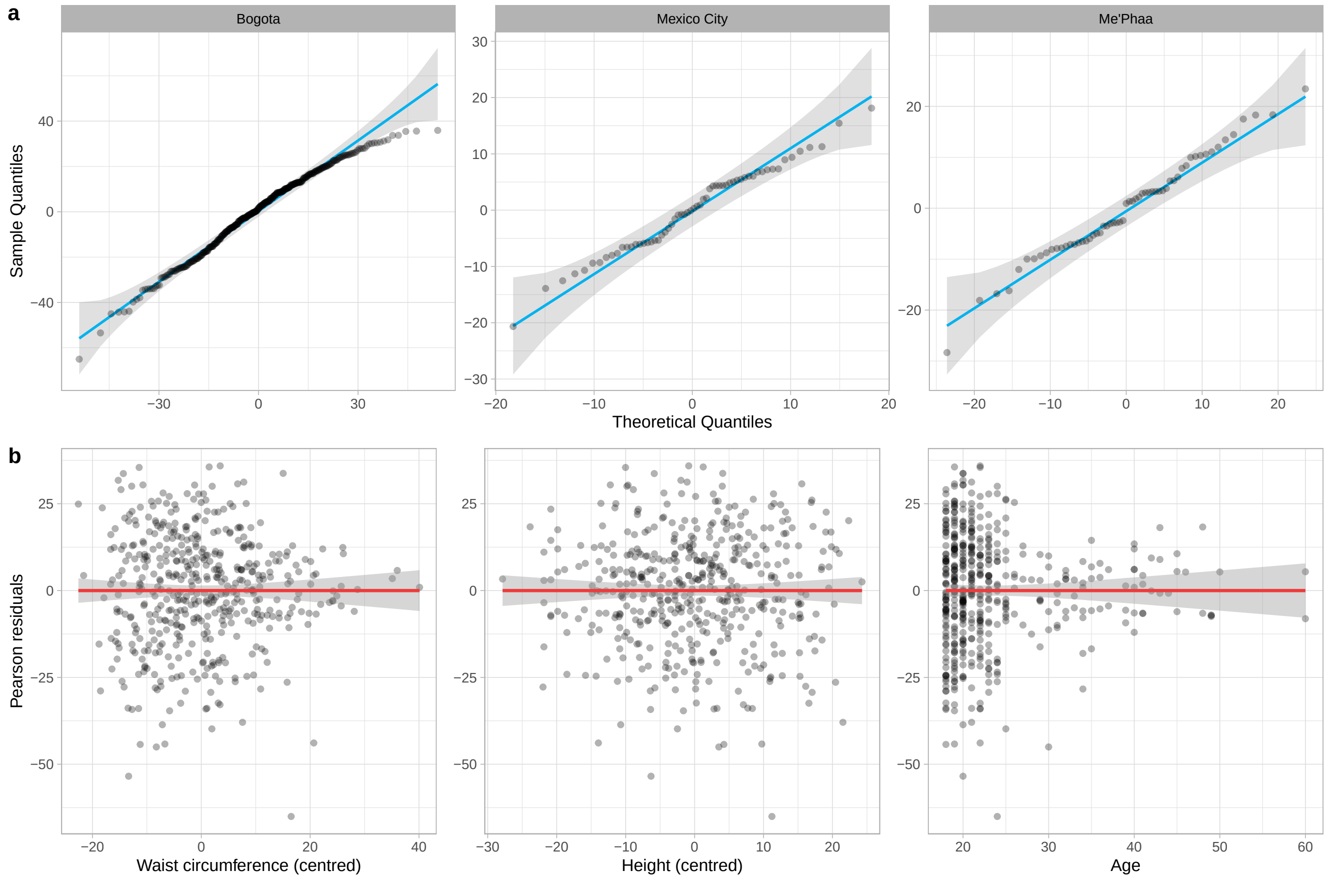

**Figure S2. Model diagnostics.** (**a**) Residual distribution for each sample. (**b**) Linearity in each (single term) fixed factor.

##### 3.3.2 Table 4. Model 3 summary, including Height × Waist Interaction

```
slopes <- sim_slopes(mod3,
           pred = Height_c,
           modx = Waist_c)

slo <- slopes$slopes
slo$CI <- paste(round(slo$`2.5%`, 3),
                round(slo$`97.5%`, 3), sep = " - ")
slo <- slo[,c(1:3,8,6:7)]

rownames(slo) <- c("WC(c) - 1 SD  = ",
                   "WC(c) Mean  = ",
                   "WC(c) + 1 SD  = ")

rownames(slo) <- paste0(rownames(slo),
                        round(slo$`Value of Waist_c`, 2))
slo <- summaSig(slo, 6)
slo[,1] <- NULL
colnames(slo) <- colnames(tabm2)
mod3Tab <- rbind(tabm3, slo)
Tab4 <- kable(mod3Tab,
      align = "c",
      caption = "\\textbf{Table 4.} Model 3 summary, including
```

```
    Height $\\times$ Waist Interaction",
    booktabs = TRUE,
    escape = FALSE) %>%
  pack_rows(group_label = "Simple slope analysis for H(c) at different values of WC(c)",
            start_row = 9,
            end_row = 11,
            hline_after = TRUE,
            bold = FALSE) %>%
  kable_styling(latex_options = "HOLD_position") %>%
  footnote(general = paste0("As waist reference, the centred values used
                          are equivalent to -1 SD (",
                          round(mean(data$Waist) - sd(data$Waist), 2),
                          " cm), mean (",
                          round(mean(data$Waist), 2),
                          " cm), and +1 SD (",
                          round(mean(data$Waist) + sd(data$Waist), 2),
                          " cm). Women and Bogota were used as reference
                          categories for Sex and Sample, respectively. For model terms:
                          WC(c) = Waist circumference (centred); H(c) = Height
                          (centred); S = Sample. Significant predictors are in bold."),
    escape = FALSE,
    threeparttable = TRUE)
Tab4
```

**Table 4.** Model 3 summary, including Height × Waist Interaction

| | *B* | *SE*(*B*) | 95% CI | *t* | *p* |
|---|---|---|---|---|---|
| (Intercept) | 61.40 | 3.77 | 53.983 — 68.815 | 16.27 | **<0.0001** |
| Age | 0.17 | 0.16 | -0.153 — 0.495 | 1.04 | 0.299 |
| H(c) | 0.16 | 0.12 | -0.077 — 0.403 | 1.33 | 0.183 |
| S(Mexico City) | -9.32 | 2.87 | -14.96 — -3.681 | -3.25 | **0.001** |
| S(Me'Phaa) | -16.78 | 3.53 | -23.722 — -9.83 | -4.75 | **<0.0001** |
| Sex(men) | 6.01 | 2.26 | 1.564 — 10.458 | 2.66 | **0.008** |
| WC(c) | -0.28 | 0.09 | -0.46 — -0.11 | -3.20 | **0.001** |
| WC(c) × H(c) | -0.02 | 0.01 | -0.036 — -0.003 | -2.30 | **0.022** |
| Simple slope analysis for H(c) at different values of WC(c) | | | | | |
| WC(c) - 1 SD = -10.4 | 0.37 | 0.15 | 0.075 — 0.657 | 2.47 | **0.014** |
| WC(c) Mean = 0 | 0.16 | 0.12 | -0.077 — 0.403 | 1.33 | 0.183 |
| WC(c) + 1 SD = 10.4 | -0.04 | 0.15 | -0.341 — 0.261 | -0.26 | 0.795 |

*Note:*
As waist reference, the centred values used are equivalent to -1 SD (67.49 cm), mean (77.89 cm), and +1 SD (88.3 cm). Women and Bogota were used as reference categories for Sex and Sample, respectively. For model terms: WC(c) = Waist circumference (centred); H(c) = Height (centred); S = Sample. Significant predictors are in bold.

##### 3.3.3 Figure 3. Model 3 estimates and Height × Waist Interaction

```
Fig3a <- plot_summs(
  mod3,
  coefs = c("Age" = "Age",
            "H(c)" = "Height_c",
            "S(Mexico City)" = "SampleMexico City",
            "S(Me'Phaa)" = "SampleMe'Phaa",
            "Sex(Men)" = "SexMen",
            "WC(c)" = "Waist_c",
            "H(c) × WC(c)" = "Height_c:Waist_c")) +
  theme_light() +
  ylab("Terms")

Fig3b <- interact_plot(
  mod3,
  pred = Height_c,
  modx = Waist_c,
  interval = TRUE,
  legend.main = "WC(c) \n reference",
              colors = colfunc(3)) +
  theme_light() +
  ylab("Self-reported health") +
  xlab("Height (c)")

Fig3cDat <- johnson_neyman(
  mod3,
  pred = Height_c,
  modx = Waist_c,
  alpha = .05,
  sig.color = colfunc(2)[1],
  insig.color = colfunc(2)[2])

Fig3c <- Fig3cDat$plot +
  theme_light() +
  labs(title = NULL) +
  ylab("Slope of Height (c)") +
  xlab("Waist circumference (c)") +
  labs(fill = "Significance")

Fig3 <- ggarrange(Fig3a,
                  ggarrange(Fig3b,
                            Fig3c,
                            nrow = 2,
                            align = "v",
                            labels = c("b", "c")),
                  ncol = 2,
                  labels = "a")
Fig3
```

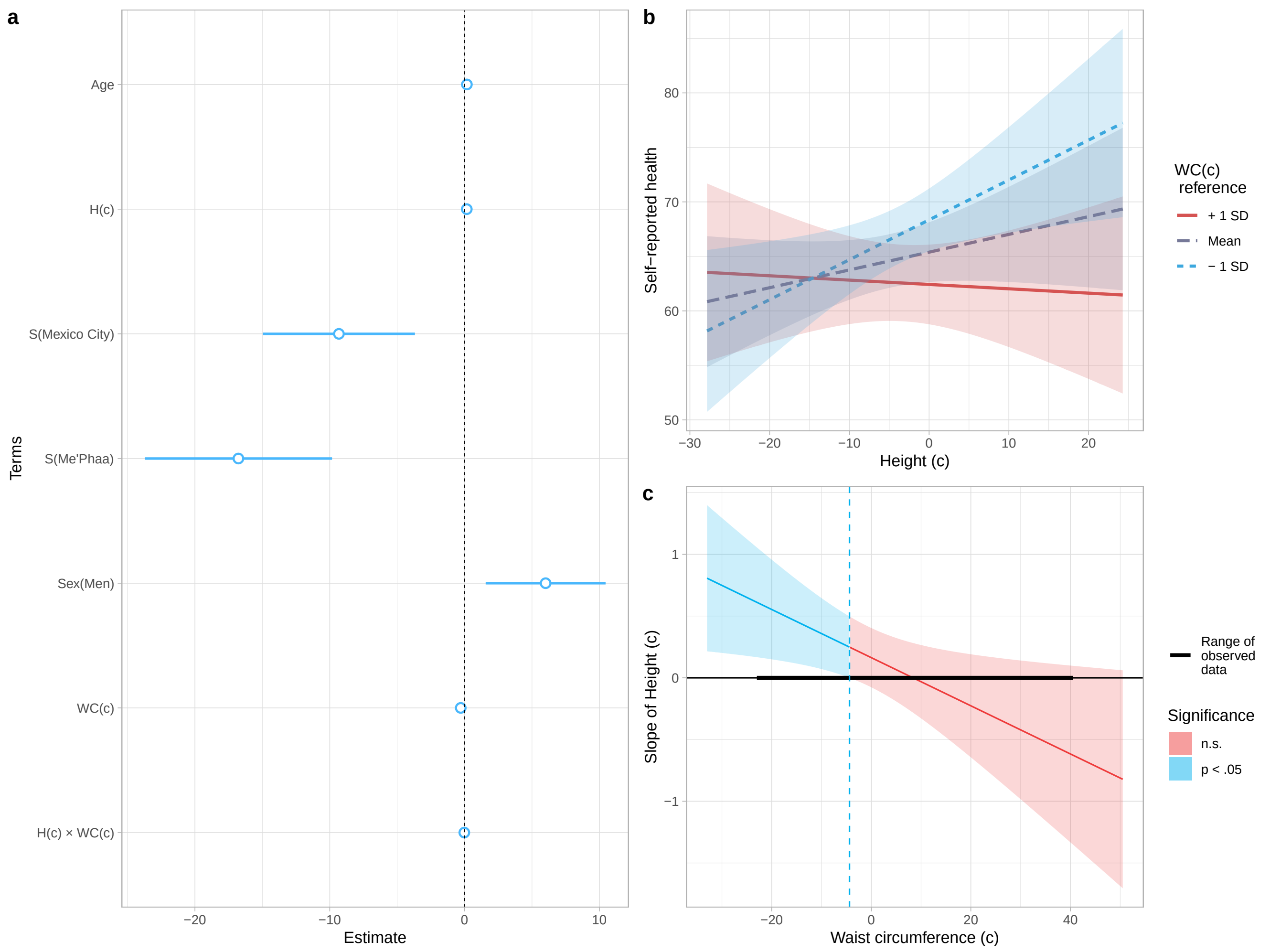

**Figure 3. Model 3 estimates and interaction between Height and Waist.** Values of Height and Waist were centred: for Height, uncentred mean ± SD = 163.85 ± 9.86 cm; for Waist circumference, uncentred mean ± SD = 77.89 ± 10.4 cm. (**a**) Estimates and 95% CI for each model term. For categorical predictors, women and Bogota were used as reference levels. For model terms, WC(c) = Waist circumference (centred); H(c) = Height (centred); S = Sample. (**b**) Interaction between Height and Waist. As waist reference, -1 SD (67.49 cm), mean (77.89 cm), and +1 SD (88.3 cm) values were used, showed on a blue to red colour scale. (**c**) Johnson-Neyman plot, showing for which values of Waist (centred), the slope of Height (centred) is significant as a predictor of Self-reported health; these slopes are predicted to be significant and positive for centred Waists circumferences below -4.38 (73.51 cm), and negative above 68.85 (146.74 cm; not shown as it is a prediction for extreme values, beyond the ones found in any of our samples).

#### 3.4 Alternative final model (Model 3A)

Given the extensive literature showing visceral fat to be a reliable marker of abdominal adiposity, and its health-related costs, we created an alternative version of the final model (Model 3), by replicating the model selection process, but replacing each instance of `Heigth` (centred) for `VisceralFat` (centred). This is, fitting an alternative Model 2, and repeating the same selection process.

##### 3.4.1 Table S9. Model 3A summary

```
# Fit model
data$VisFat_c <- c(scale(data$VisceralFat, scale = FALSE))
mod1a <- lm(Health ~
            Sex * VisFat_c * Height_c * Sample + Age +
```

```
             Hip + Fat + VisFat_c + Weight + Muscle + BMI,
           data = data)
mod2a <- lm(Health ~
             Sex * VisFat_c * Height_c * Sample + Age,
           data = mod1a$model,
           na.action = "na.fail")
# Selection process
fitta <- dredge(mod2a)
options(digits = 2)
m.sel <- model.sel(fitta)
# Select best model and assign it as Model 3A
mod3a <- get.models(fitta, subset = 1)[[1]]

# Create and format summary data.frame
s.mod3a <- summary(mod3a)
ci.mod3a <- as.data.frame(confint(mod3a))
ci.mod3a$CI <- paste(round(ci.mod3a$`2.5 %`, 3), round(ci.mod3a$`97.5 %`, 3), sep = " - ")
tabm3a <- as.data.frame(s.mod3a$coefficients)
tabm3a <- cbind(tabm3a, ci.mod3a$CI)
tabm3a <- summaSig(tabm3a, 4)
tabm3a <- summTerms(tabm3a)
row.names(tabm3a) <- str_replace(row.names(tabm3a),
                                    "VisFat_c", "VF(c)")
tabm3a <- tabm3a[,c(1,2,5,3,4)]
colnames(tabm3a) <- summCols
# Simple slope analysis
slopesa <- sim_slopes(mod3a,
           pred = Height_c,
           modx = VisFat_c)
sloa <- slopesa$slopes
sloa$CI <- paste(round(sloa$`2.5%`, 3),
                round(sloa$`97.5%`, 3), sep = " - ")
sloa <- sloa[,c(1:3,8,6:7)]
rownames(sloa) <- c("VF(c) - 1 SD  = ",
                   "VF(c) Mean  = ",
                   "VF(c) + 1 SD  = ")
rownames(sloa) <- paste0(rownames(sloa),
                        round(sloa$`Value of VisFat_c`, 2))
sloa <- summaSig(sloa, 6)
sloa[,1] <- NULL
colnames(sloa) <- colnames(tabm2)
mod3aTab <- rbind(tabm3a, sloa)
# Final table
TabS9 <- kable(mod3aTab,
      align = "c",
      caption = "\\textbf{Table S9.} Model 3A summary, including
      Height $\\times$ Visceral Fat Interaction",
      booktabs = TRUE,
      escape = FALSE) %>%
  pack_rows(group_label = "Simple slope analysis for H(c) at different values of VF(c)",
            start_row = 11,
            end_row = 13,
            hline_after = TRUE,
            bold = FALSE) %>%
  kable_styling(latex_options = "HOLD_position") %>%
  footnote(general = paste0("$R^2$ = ", round(s.mod3a$r.squared, 3),
```

```
    ", $R^2_{adjusted}$ = ",
    round(s.mod3a$adj.r.squared, 3), ", $F$(",
    paste(s.mod3a$fstatistic[2],
          s.mod3a$fstatistic[3], sep = ", "),
    ") = ", round(s.mod3a$fstatistic[1], 2), ", $p$ ",
    pvalr(pf(s.mod3a$fstatistic[1],
             s.mod3a$fstatistic[2],
             s.mod3a$fstatistic[3],
             lower.tail = FALSE),
          digits = 4),
    ". Women and Bogota were used as reference categories for Sex and Sample,
    respectively. For model terms: WC(c) = Waist circumference (centred); H(c) =
    Height (centred); S = Sample. Significant predictors are in bold. As visceral
    fat reference, the centred values used are equivalent to -1 SD (",
                            round(mean(data$VisceralFat) -
                                    sd(data$VisceralFat), 2),
                            "), mean (", round(mean(data$VisceralFat), 2),
                            "), and +1 SD (", round(mean(data$VisceralFat) +
                                    sd(data$VisceralFat), 2),
                            "). Women and Bogota were used as reference
                            categories for Sex and Sample, respectively. For model terms:
                            WC(c) = Waist circumference (centred); VF(c) = Visceral Fat
                            (centred); S = Sample. Significant predictors are in bold."),
    escape = FALSE,
    threeparttable = TRUE)
TabS9
```

**Table S9.** Model 3A summary, including Height × Visceral Fat Interaction

| | $B$ | $SE(B)$ | 95% CI | $t$ | $p$ |
|---|---|---|---|---|---|
| (Intercept) | 62.52 | 1.76 | 59.067 — 65.976 | 35.56 | **<0.0001** |
| H(c) | 0.05 | 0.12 | -0.19 — 0.297 | 0.43 | 0.667 |
| S(Mexico City) | -4.68 | 3.72 | -11.982 — 2.632 | -1.26 | 0.209 |
| S(Me'Phaa) | -10.40 | 4.23 | -18.715 — -2.081 | -2.46 | **0.014** |
| Sex(men) | 9.39 | 2.70 | 4.078 — 14.702 | 3.47 | **<0.001** |
| VF(c) | -2.20 | 0.79 | -3.745 — -0.65 | -2.79 | **0.005** |
| H(c) × VF(c) | -0.10 | 0.04 | -0.173 — -0.028 | -2.73 | **0.007** |
| S(Mexico City) × Sex(men) | -6.74 | 4.94 | -16.459 — 2.969 | -1.36 | 0.173 |
| S(Me'Phaa) × Sex(men) | -10.91 | 5.28 | -21.297 — -0.527 | -2.06 | **0.039** |
| Sex(men) × VF(c) | 1.97 | 0.91 | 0.182 — 3.766 | 2.16 | **0.031** |
| Simple slope analysis for H(c) at different values of VF(c) | | | | | |
| VF(c) - 1 SD = -2.88 | 0.34 | 0.16 | 0.037 — 0.649 | 2.20 | **0.028** |
| VF(c) Mean = 0 | 0.05 | 0.12 | -0.19 — 0.297 | 0.43 | 0.667 |
| VF(c) + 1 SD = 2.88 | -0.24 | 0.17 | -0.571 — 0.099 | -1.38 | 0.167 |

*Note:*
$R^2 = 0.188$, $R^2_{adjusted} = 0.172$, $F(9, 463) = 11.87$, $p < 0.001$. Women and Bogota were used as reference categories for Sex and Sample, respectively. For model terms: WC(c) = Waist circumference (centred); H(c) = Height (centred); S = Sample. Significant predictors are in bold. As visceral fat reference, the centred values used are equivalent to -1 SD (2.39), mean (5.27), and +1 SD (8.15). Women and Bogota were used as reference categories for Sex and Sample, respectively. For model terms: WC(c) = Waist circumference (centred); VF(c) = Visceral Fat (centred); S = Sample. Significant predictors are in bold.

##### 3.4.2 Figure S3. Model 3 estimates and Height × Waist Interaction

```
FigS3a <- plot_summs(
  mod3a,
  coefs = c("H(c)" = "Height_c",
            "S(Mexico City)" = "SampleMexico City",
            "S(Me'Phaa)" = "SampleMe'Phaa",
            "Sex(Men)" = "SexMen",
            "VF(c)" = "VisFat_c",
            "H(c) × VF(c)" = "Height_c:VisFat_c",
            "S(Mexico City) × Sex(Men)" = "SampleMexico City:SexMen",
            "S(Me'Phaa) × Sex(Men)" = "SampleMe'Phaa:SexMen",
            "Sex(Men) × VF(c)" = "SexMen:VisFat_c")) +
  theme_light() +
  ylab("Terms")

FigS3b <- interact_plot(
  mod3a,
  pred = Height_c,
  modx = VisFat_c,
  interval = TRUE,
  legend.main = "VF(c) \n reference",
              colors = colfunc(3)) +
  theme_light() +
  ylab("Self-reported health") +
  xlab("Height (c)")

FigS3dDat <- johnson_neyman(
  mod3a,
  pred = Height_c,
  modx = VisFat_c,
  alpha = .05,
  sig.color = colfunc(2)[1],
  insig.color = colfunc(2)[2])

FigS3c <- FigS3dDat$plot +
  theme_light() +
  labs(title = NULL) +
  ylab("Slope of Height (c)") +
  xlab("Visceral Fat (c)") +
  labs(fill = "Significance")

FigS3 <- ggarrange(FigS3a,
                   ggarrange(FigS3b,
                             FigS3c,
                             nrow = 2,
                             align = "v",
                             labels = c("b", "c")),
                   ncol = 2,
                   labels = "a")
FigS3
```

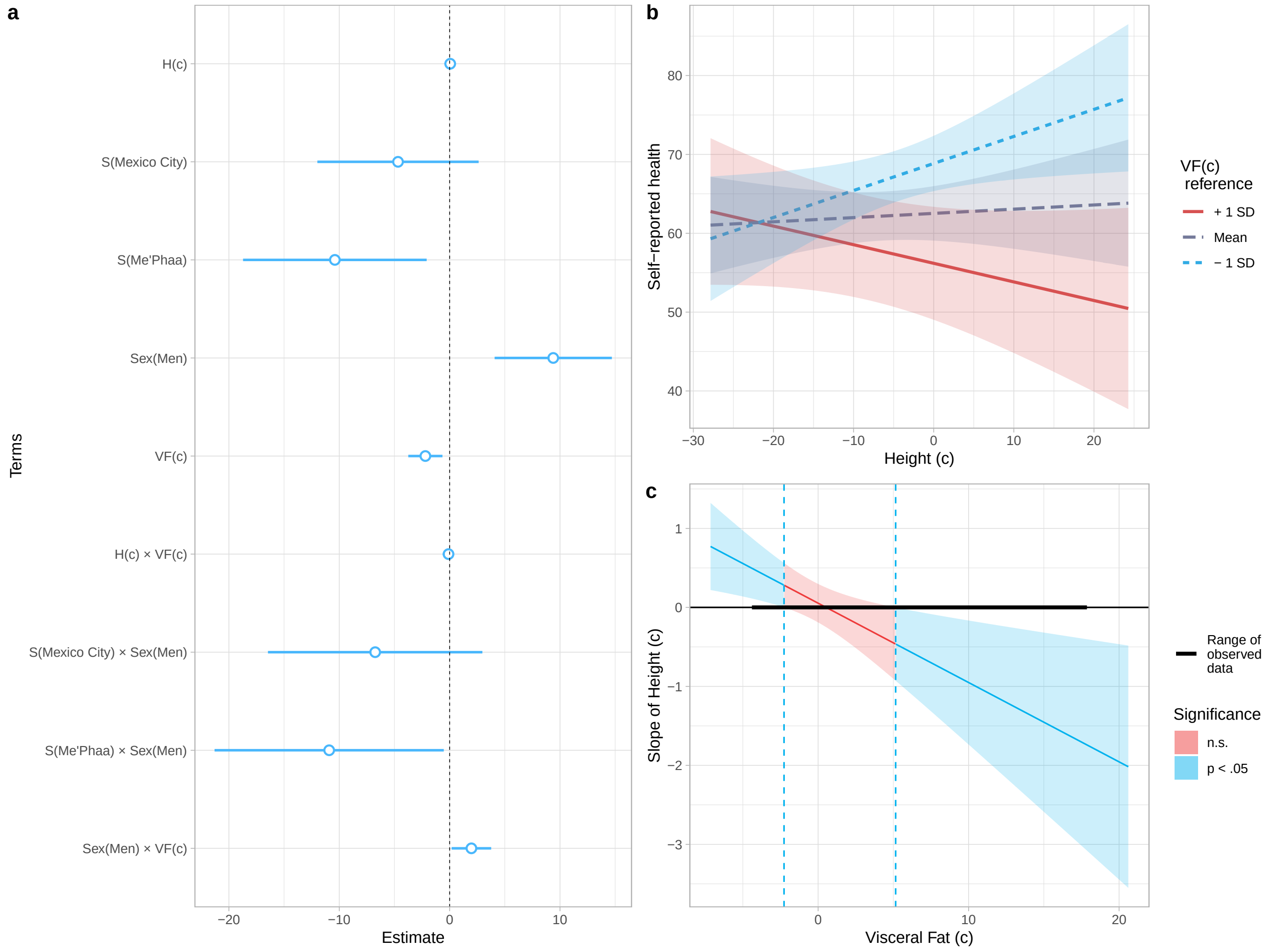

**Figure S3. Model 3A estimates and interactions.** Values of Height and Visceral Fat were centred: for Height, uncentred mean ± SD = 163.85 ± 9.86 cm; for Visceral Fat, uncentred mean ± SD = 5.27 ± 2.88. (**a**) Estimates and 95% CI for each model term. For categorical predictors, women and Bogota were used as reference levels. For model terms, VF(c) = Visceral Fat (centred); H(c) = Height (centred); S = Sample. (**b**) Interaction between Height and Visceral Fat. As Visceral Fat reference, -1 SD (2.39), mean (5.27), and +1 SD (8.15) values were used, showed on a blue to red colour scale. (**c**) Johnson-Neyman plot, showing for which values of Visceral Fat (centred), the slope of Height (centred) is significant as a predictor of Self-reported health; these slopes are predicted to be significant and positive for centred Visceral Fat levels below -2.27 (3.002 uncentred), and negative for values above 5.15 (10.42 uncentred).

##### 3.4.3 Table S10. Information criteria for the alternative and final models

In addition to being more parsimonious, Model 3 (mod3) had a lower Akaike information criterion ($AICc$), and higher Akaike weight ($w_i(AICc)$) than the alternative final model, but the $\Delta AICc$ was less that 2 units.

```
aicf1_2 <- AICctab(mod3,
                   mod3a,
                   weights = TRUE,
                   base = TRUE)
class(aicf1_2) <- "data.frame"
tabs10 <- aicf1_2
row.names(tabs10) <- c("Model 3", "Alternative Model 3")
tabs10$weight <- format(round(tabs10$weight, 3))

TabS10 <- kable(
```

```
  tabs10,
  align = "c",
  digits = 20,
  caption = "\\textbf{Table S10.} Information criteria for alternative and final models",
  col.names = c("$AICc$", "$\\Delta AICc$", "$df$", "$w_{i}(AICc)$"),
  booktabs = TRUE,
  escape = FALSE) %>%
  kable_styling(latex_options = "HOLD_position") %>%
  footnote(general = paste0("Model 3 is close to ",
                            format(round(aicf1_2[1,4]/aicf1_2[2,4], 10),
                                   big.mark = ","),
                            " times more likely to be the best model
                            compared to the Aternative Model 3.
                            For a detailed description of values,
                            see \\\\href{https://www.shorturl.at/iGIKT}{ICtab}
                            function documentation."),
           escape = FALSE,
           threeparttable = TRUE)
TabS10
```

**Table S10.** Information criteria for alternative and final models

| | $AICc$ | $\Delta AICc$ | $df$ | $w_i(AICc)$ |
|---|---|---|---|---|
| Model 3 | 3999 | 0.0 | 9 | 0.68 |
| Alternative Model 3 | 4001 | 1.5 | 11 | 0.32 |

*Note:*
Model 3 is close to 2.1 times more likely to be the best model compared to the Aternative Model 3. For a detailed description of values, see ICtab function documentation.

### 4 Session info (for reproducibility)

```
library(pander)
pander(sessionInfo(), locale = FALSE)
```

**R version 3.6.1 (2019-07-05)**

**Platform:** x86_64-w64-mingw32/x64 (64-bit)

**attached base packages:** *stats4*, *stats*, *graphics*, *grDevices*, *utils*, *datasets*, *methods* and *base*

**other attached packages:** *pander(v.0.6.3)*, *lmSupport(v.2.9.13)*, *interactions(v.1.1.1)*, *qqplotr(v.0.0.3)*, *MuMIn(v.1.43.6)*, *rstatix(v.0.2.0)*, *data.table(v.1.12.6)*, *lemon(v.0.4.3)*, *bbmle(v.1.0.20)*, *Hmisc(v.4.2-0)*, *Formula(v.1.2-3)*, *lattice(v.0.20-38)*, *jtools(v.2.0.1)*, *survival(v.2.44-1.1)*, *car(v.3.0-4)*, *carData(v.3.0-2)*, *ryouready(v.0.4)*, *kableExtra(v.1.1.0)*, *psych(v.1.8.12)*, *ggpubr(v.0.2.3)*, *magrittr(v.1.5)*, *gridExtra(v.2.3)*, *forcats(v.0.4.0)*, *stringr(v.1.4.0)*, *dplyr(v.0.8.3)*, *purrr(v.0.3.3)*, *readr(v.1.3.1)*, *tidyr(v.1.0.0)*, *tibble(v.2.1.3)*, *ggplot2(v.3.2.1)*, *tidyverse(v.1.2.1)*, *osfr(v.0.2.4)* and *knitr(v.1.25)*

**loaded via a namespace (and not attached):** *readxl(v.1.3.1)*, *backports(v.1.1.5)*, *VGAM(v.1.1-1)*, *plyr(v.1.8.4)*, *lazyeval(v.0.2.2)*, *sp(v.1.3-1)*, *splines(v.3.6.1)*, *unmarked(v.0.12-3)*, *urltools(v.1.7.3)*, *digest(v.0.6.22)*, *htmltools(v.0.4.0)*, *gdata(v.2.18.0)*, *checkmate(v.1.9.4)*, *AICcmodavg(v.2.2-2)*, *cluster(v.2.1.0)*, *openxlsx(v.4.1.2)*, *modelr(v.0.1.5)*, *sandwich(v.2.5-1)*, *colorspace(v.1.4-1)*, *rvest(v.0.3.4)*, *haven(v.2.1.1)*, *xfun(v.0.10)*, *crayon(v.1.3.4)*, *jsonlite(v.1.6)*, *lme4(v.1.1-21)*, *zeallot(v.0.1.0)*, *zoo(v.1.8-6)*, *glue(v.1.3.1)*, *gtable(v.0.3.0)*, *webshot(v.0.5.1)*, *DEoptimR(v.1.0-8)*, *abind(v.1.4-5)*, *scales(v.1.0.0)*, *Rcpp(v.1.0.2)*, *viridisLite(v.0.3.0)*, *xtable(v.1.8-4)*, *htmlTable(v.1.13.2)*, *ggstance(v.0.3.3)*, *foreign(v.0.8-71)*, *htmlwidgets(v.1.5.1)*, *httr(v.1.4.1)*, *gplots(v.3.0.1.1)*, *RColorBrewer(v.1.1-2)*, *acepack(v.1.4.1)*, *pkgconfig(v.2.0.3)*, *nnet(v.7.3-12)*, *crul(v.0.8.4)*, *tidyselect(v.0.2.5)*, *labeling(v.0.3)*, *rlang(v.0.4.1)*, *reshape2(v.1.4.3)*, *munsell(v.0.5.0)*, *cellranger(v.1.1.0)*, *tools(v.3.6.1)*, *cli(v.1.1.0)*, *generics(v.0.0.2)*, *broom(v.0.5.2)*, *evaluate(v.0.14)*, *yaml(v.2.2.0)*, *fs(v.1.3.1)*, *zip(v.2.0.4)*, *robustbase(v.0.93-5)*, *caTools(v.1.17.1.2)*, *nlme(v.3.1-140)*, *xml2(v.1.2.2)*, *compiler(v.3.6.1)*, *pbkrtest(v.0.4-7)*, *rstudioapi(v.0.10)*, *curl(v.4.2)*, *ggsignif(v.0.6.0)*, *stringi(v.1.4.3)*, *highr(v.0.8)*, *Matrix(v.1.2-17)*, *nloptr(v.1.2.1)*, *vctrs(v.0.2.0)*, *pillar(v.1.4.2)*, *lifecycle(v.0.1.0)*, *triebeard(v.0.3.0)*, *pwr(v.1.2-2)*, *cowplot(v.1.0.0)*, *bitops(v.1.0-6)*, *raster(v.3.0-7)*, *R6(v.2.4.0)*, *latticeExtra(v.0.6-28)*, *KernSmooth(v.2.23-15)*, *rio(v.0.5.16)*, *codetools(v.0.2-16)*, *boot(v.1.3-22)*, *MASS(v.7.3-51.4)*, *gtools(v.3.8.1)*, *assertthat(v.0.2.1)*, *withr(v.2.1.2)*, *httpcode(v.0.2.0)*, *mnormt(v.1.5-5)*, *parallel(v.3.6.1)*, *hms(v.0.5.2)*, *grid(v.3.6.1)*, *rpart(v.4.1-15)*, *minqa(v.1.2.4)*, *rmarkdown(v.1.16)*, *numDeriv(v.2016.8-1.1)*, *gvlma(v.1.0.0.3)*, *lubridate(v.1.7.4)* and *base64enc(v.0.1-3)*

### 5 Supplementary Reference

1. Lugo A, L. H., García E, H. I. & Gómez R, C. Confiabilidad del cuestionario de calidad de vida en salud SF-36 en Medellín, Colombia. *Rev. Fac. Nac. Salud Publica* **24**, 37–50, http://www.scielo.org.co/scielo.php?script=sci_arttext&pid=S0120-386X2006000200005 (2006).
